# Age-dependent mechanical stress underlies unexpected vestibular differentiation and regeneration patterns

**DOI:** 10.64898/2026.11.05.764569

**Authors:** Shahar Inbar Kasirer, Michal Shraga, Tim Dullweber, Olga Loza, Anna Erzberger, David Sprinzak

## Abstract

Loss of inner-ear sensory hair cells (HCs) is a major cause of hearing and balance disorders. Promoting HC regeneration offers a potential therapeutic strategy, but regenerative capacity is limited in adult mammals and declines rapidly during early postnatal development in mice. HCs can regenerate through trans-differentiation of neighboring supporting cells in a process controlled by Notch mediated lateral inhibition. However, the factors underlying the decrease in regeneration capacity are poorly understood. Here, we combine live imaging with single-HC ablation in mouse vestibular explants to track the dynamics of HC development and regeneration. We reveal unexpected age-dependent deviations from the classical lateral inhibition model, in which new HCs differentiate adjacent to existing HCs. We find that mechanical stress changes with age and that perturbing it alters the observed differentiation patterns. A mathematical model incorporating local mechanical regulation of Notch signaling captures both HC differentiation and regeneration patterns across ages. Together, these findings identify mechanical stress as a crucial regulator of HC differentiation and support an underlying mechanism in which the interplay between local mechanics and Notch signaling shapes regenerative responses. This coupling offers a framework for understanding the age-dependent decline in HC regeneration and motivates investigation of mechanical regulation as a potential therapeutic target.

## Introduction

Loss of sensory hair cells (HCs) in the inner ear is considered a major cause of hearing and balance disorders in human patients^1–3^. A potential treatment for hearing and balance disorders is the promotion of HC regeneration^4–6^. For mammals, the regeneration capacity declines rapidly at early postnatal stages (approximately after postnatal day 14 in mice^7^). This ability to regenerate is more prominent in the vestibular system, which retains a limited regeneration capacity until adulthood^5,8^. However, what controls regeneration capacity and what leads to its decline at postnatal stages is not clear.

The vestibular system is composed of five sensory organs – the utricle and saccule, sensing linear acceleration and head orientation, and three cristae, sensing head rotations^9^ (Figure 1a). Each of these sensory organs is composed of sensory HCs interspersed with non-sensory supporting cells (SCs)^10^. It is known that in neonatal mice, HC regeneration occurs through proliferation and direct trans-differentiation of SCs^11,12^. In adults, HC regeneration predominantly occurs through trans-differentiation without proliferation^13^.

**Figure 1.**
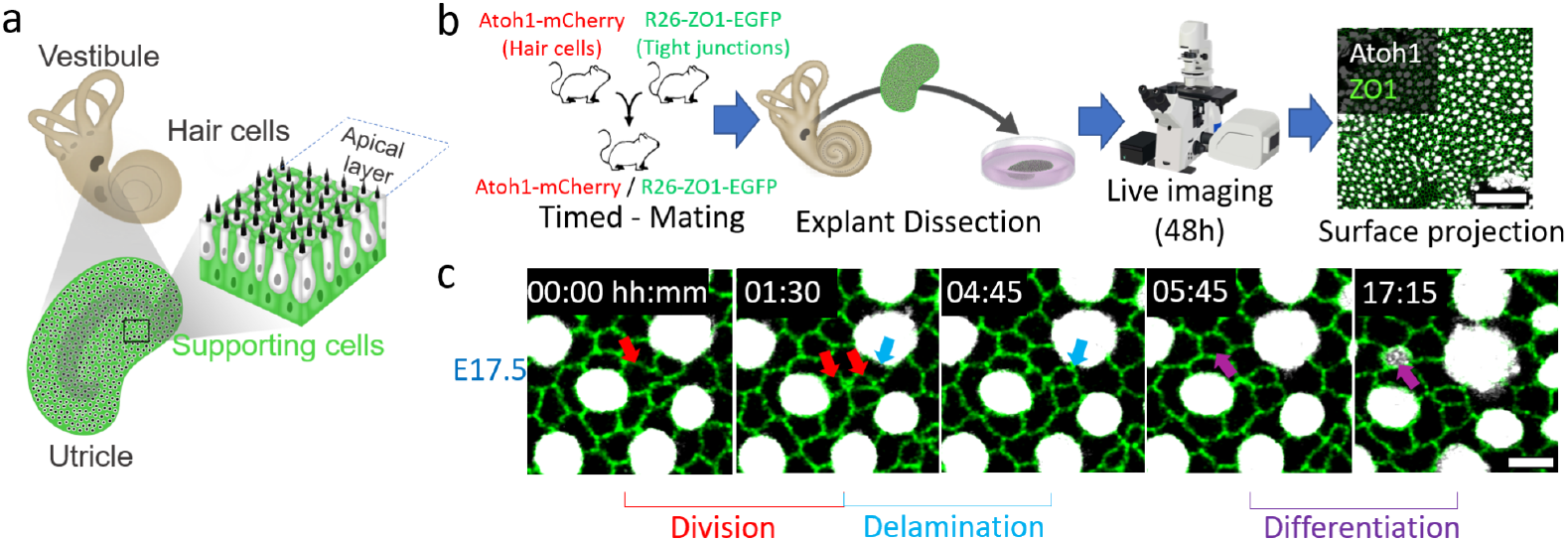
Live imaging of mouse utricle explants enables tracking developmental dynamics. **(a)** Schematic illustration of the utricle, one of the sensory organs in the vestibular system within the inner ear. It is composed of sensory HCs (white) and non-sensory supporting cells (green). **(b)** Live imaging assay starts with a timed mating between transgenic mice from two reporter lines. Utricle explants from embryos/neonates are dissected at the desired stage and live imaged for 48 hours, 4 explants at a time. The apical layer is extracted from the resulting confocal stacks via surface projection. Scale bar - 50μm. **(c)** A filmstrip from a live imaging of an embryonic day 17.5 (E17.5) utricle explant, demonstrating cell division (red arrows), delamination (blue arrows) and differentiation (magenta arrows) events. Time from movie start is indicated. Scale bar - 5μm. HCs (white) are marked by Atoh1-mCherry, apical cell boundaries (green) are marked by ZO1-EGFP (b-c).

The main pathway that regulates the differentiation of prosensory cells into HCs and SCs is the Notch pathway^10^. In the commonly used model, differentiation is achieved through the process of lateral inhibition (LI), mediated by Notch signaling^14^. During this process, HCs express high levels of Notch ligands, promoting the activation of Notch signaling in neighboring cells. This activation prevents neighboring cells from becoming HCs themselves. This results in the prototypical salt-and-pepper pattern, where each HC is surrounded by SC neighbors^15^. This model was tested in the mouse vestibular system by blocking Notch signaling, which resulted in a significant increase in HC numbers^16,17^. Importantly, downregulation of Notch signaling enhances HC regeneration capacity following HC damage induced by ototoxic drugs or genetic manipulation^16–18^. However, the regenerative response to Notch downregulation is weaker for adults compared with young mice^19,20^. Therefore, Notch inhibition alone is not sufficient as HC regeneration treatment.

HC regeneration has also been suggested to be regulated by mechanical forces within the sensory domains. A previously postulated hypothesis, termed “the parking brake hypothesis”, suggests that higher mechanical stiffness of the tissue leads to reduced HC regeneration with age^21^. Supporting evidence for this hypothesis includes the observations of differences in tissue stiffness between the avian inner ear^22^, where regeneration persists into adulthood, and the mammalian inner ear. Similar mechanical differences were observed between early and late postnatal stages in mice^22^. This difference was also correlated with enhancement of F-actin bands at the later postnatal stage^22,23^. Finally, recent work in inner ear organoids showed that HC differentiation is regulated by the stiffness of extracellular matrix^24^. In light of these observations, mechanical regulation of HC regeneration seems plausible. However, the mechanistic details of this mechano-regulation are unclear. Specifically, it is unknown how the interplay between mechanics and Notch-mediated LI locally coordinates HC differentiation and regeneration.

In this study, we use live imaging of mouse vestibular explants to track the dynamics of HC differentiation and regeneration during late embryonic and early postnatal stages. Quantitative analysis of differentiation and regeneration dynamics reveals age-dependent differences in local differentiation patterns. We find that HCs differentiate next to existing HCs, in contrast to the expectation from the LI model. This deviation from the classical model is more prominent at later stages, where regeneration capacity is more limited. We further show that these age-dependent differentiation patterns are correlated with changes in mechanical stress and are affected by reducing tissue contractility. Finally, we use mathematical modeling to show that the interplay between local mechanical stress and LI can explain the observed patterns. Our findings implicate the interplay between mechanical stress and Notch signaling as an important factor underlying age-dependent decline in regenerative capacity.

## Results

### Live imaging of utricle explants enables tracking developmental dynamics

To investigate local HC differentiation dynamics we established a live imaging assay in mouse vestibular explants from a double transgenic mouse (Figure 1b). This assay is based on our previous cochlear explant assay^25,26^, which allows tracking apical cell morphology using a transgenic mouse line containing a ZO1-EGFP tight junction marker localized to the apical boundaries^27^. In addition, to mark HC differentiation, we used an Atoh1-mCherry mouse line that reports early HC differentiation^28,29^. We chose to focus on the utricle, the linear horizontal acceleration sensor, which has been commonly used for explant protocols in previous works^11,30,31^. Explants were extracted from timed-mated pregnancies at the desired stage. As our goal is to study HC regeneration, we chose to compare embryonic day 17.5 (E17.5), a late embryonic stage where HC differentiation in the utricle is relatively abundant^32^, to postnatal day 0 (P0) where HC differentiation still exists, but is significantly less common^7^. These explants were imaged for 48 hours using a scanning confocal microscope. We then used a custom-made surface projection algorithm, based on the apical localization of ZO1, to extract the apical plane (Figure 1b).

The movies obtained from the live imaging of utricle explants revealed several dynamic cellular events including SC divisions, SC delaminations, and differentiation of SCs into HCs (Figure 1c, Extended Data Movie 1). We did not observe any HCs dividing, delaminating, or converting into SCs. Moreover, all events were less frequent at P0 compared to E17.5 (Extended Data Figure 1). In particular, division events were extremely rare at P0 (Extended Data Figure 1). To check that our live imaging assay exhibited similar cellular patterns as the *in vivo* ones, we analyzed the local patterns (i.e. number and type of neighbors) after 48 hours of live imaging and compared them to the *in vivo* generated patterns at the corresponding stage. The analysis showed no significant differences between the *in vivo* and *ex vivo* patterns, supporting normal differentiation behavior in the *ex vivo* assay (Extended Data Figure 2).

### Live imaging enables tracking differentiating HCs both during normal development and following induced local damage

To study differentiation dynamics, we scanned our live imaging results to detect all differentiation events. We detected multiple differentiation events at both developmental stages (Figure 2a,b). We noticed that at E17.5 the differentiating cells usually had no HC neighbors (Figure 2a, Extended Data Movie 2), as expected from LI models^15,33–35^. Interestingly, differentiating cells at P0 usually had a single HC neighbor (Figure 2b, Extended Data Movie 3), in distinct contrast to the expected behavior from LI, where HC differentiation should be inhibited by neighboring HC.

**Figure 2.**
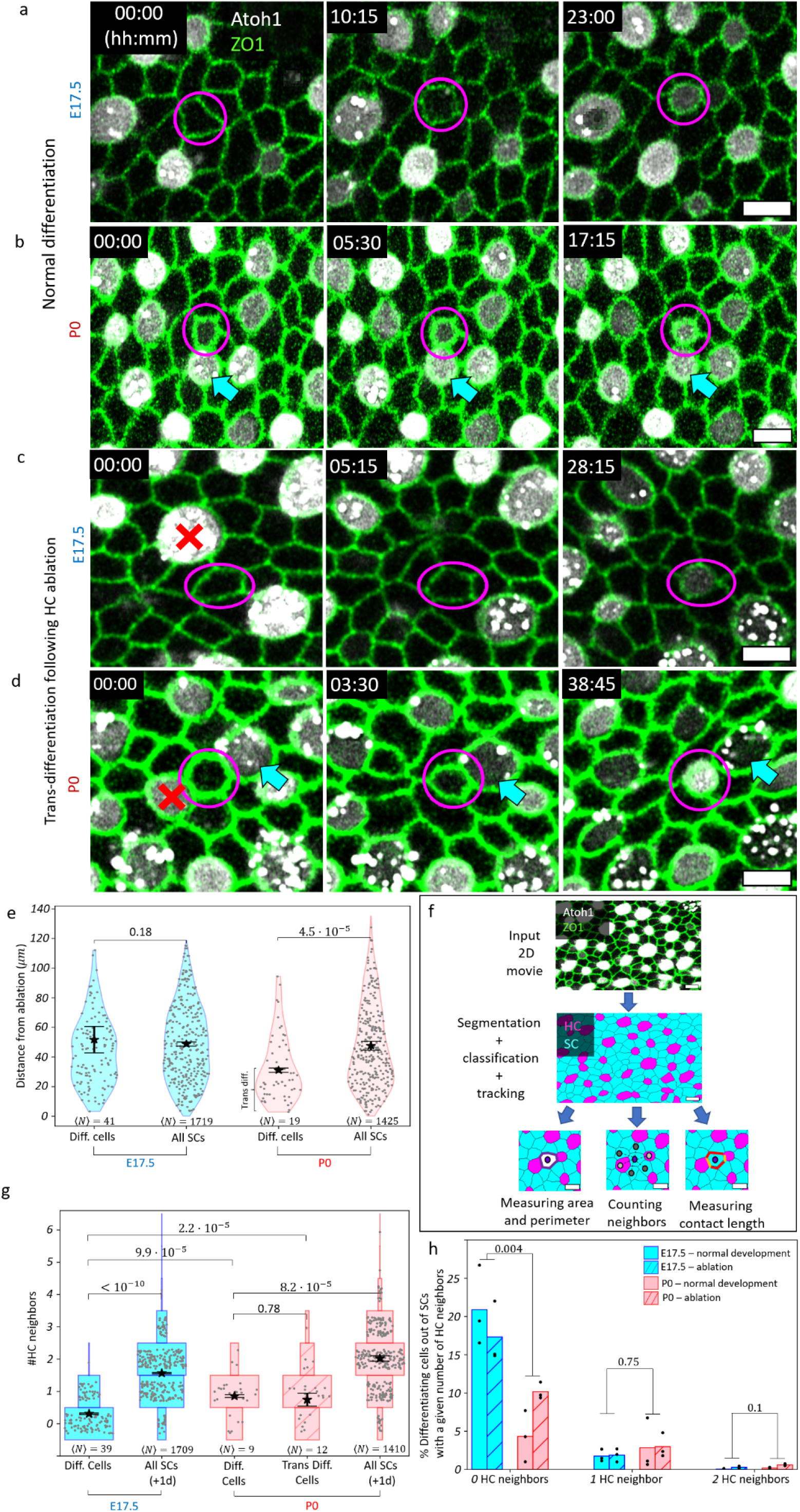
Quantitative analysis of live imaging reveals distinct differentiation and regeneration patterns for late embryonic and early postnatal stages. **(a-d)** Film strips from movies showing examples of differentiation events (purple circles, a-b) and trans-differentiation events (purple circles, c-d) following HC ablation (red X). At E17.5 (a,c), the differentiating/trans-differentiating cells typically have no HC neighbors. At P0 (b,d), the differentiating/trans-differentiating cells typically have one HC neighbor (blue arrows). **(e)** Violin plot for the distance of each differentiating cell from the ablated HC just before ablation, compared with the distance of every SC (differentiating and non-differentiating) in the same frame, for E17.5 (blue) and P0 (red). p-values were calculated by fitting a log-transformed linear mixed model where the biological repeat (three for each stage) was treated as the random effect and the group label (e.g. differentiating cells vs. all SC) as the fixed effect. **(f)** Schematic of the semi-automated quantitative analysis pipeline used. Live imaging apical surface results (after surface projection) undergo semi-automated segmentation (using ZO1 signal), classification into cell types (using Atoh1 signal), and tracking. These analyzes enable the automatic extraction of quantitative data for each cell (e.g. area, perimeter, number of neighbors and contact length with neighbors). All scale bars - 5µm (a-d,f). **(g)** Discrete violin plot (horizontal histogram) for the number of HC neighbors for each differentiating cell at E17.5 (blue) and each differentiating and trans-differentiating cell at P0 (red). The group of all SCs at the middle of each movie (after 24 hours) is used as a reference. ⟨N⟩, the average number of cells in each group (over all biological repeats), is indicated under each plot. Group means (stars) and standard errors (error bars) are indicated (e,g). p-values were calculated by fitting a generalized linear mixed model with Poisson distribution where the biological repeat (three repeats for each stage and condition) was treated as the random effect and the group as the fixed effect. **(h)** The percentage of detected differentiations out of the group of SCs with zero, one or two HC neighbors (# Differentiating cells with n HC neighbors/# SCs with n HC neighbors), for differentiating cells at E17.5 (blue) and P0 (red). Group sizes for each biological repeat were estimated using the data from the reference frame in the middle of each movie (same data reference as in g). p-values were calculated using Scheirer-Ray-Hare test (non-parametric two-way ANOVA) to compare developmental stages. Scattered data points are randomly shifted along the x-direction (e,h) and y-direction (g only). For groups with more than 100 data points only a random subset of 100 points is shown.

In addition to tracking HC differentiation during normal development, we wanted to track HC regeneration dynamics. To do that, we used a high intensity UV laser to ablate single HCs, and tracked the following dynamics using live imaging. For each explant, we ablated 3-4 distant HCs (at least 80µ*m* apart). Following ablation, we typically observed a nearby SC differentiating into an HC. The detected differentiation events exhibited a similar pattern to the one observed during normal development: Differentiating cells usually had no HC neighbor at E17.5 (Figure 2c, Extended Data Movie 4) and one HC neighbor at P0 (Figure 2d, Extended Data Movie 5).

### Single HC ablation promotes trans-differentiation of nearby SCs into HCs at P0

To check whether the observed differentiation events after HC ablation were promoted by the local damage, the distance of every SC from the nearest ablation site was measured at the initial frame, just before ablation. We then asked if there was a selection bias, namely, did SCs that were closer to an ablation site have a higher probability to differentiate? To answer this question, the distance-to-ablation for the sub-group of differentiating SCs was compared to the entire group of SCs (Figure 2e). At E17.5 there was no significant difference, i.e. no selection bias. However, at P0 we observed a high abundance of differentiating cells near the ablation site, with a peak at a distance of ∼20µ*m* (approximately 3-4 cell diameters, Figure 2e). Together with the increased number of differentiation events (Extended Data Figure 1), we concluded that local HC ablation promoted the differentiation of nearby SCs at P0. As previous studies showed that SCs replace HCs via direct trans-differentiation^11,13^, we used the average differentiation events distance-to-ablation at P0 as a threshold and marked the observed differentiations near ablation as trans-differentiation events (Figure 2e).

### Quantitative analysis reveals distinct differentiation and trans-differentiation patterns for late embryonic and early postnatal stages

As both normal differentiations (i.e. differentiations during normal development) and trans-differentiations after HC ablation exhibited the same pattern where differentiating HC had mostly no HC neighbors at E17.5 vs. one HC neighbor at P0, we decided to use quantitative analysis to further explore this behavior. We established a semi-automated analysis pipeline for segmentation and tracking of cellular events (Figure 2f). Based on this quantitative analysis, the number of apical HC neighbors (i.e. cells that share apical boundary) for each differentiating cell was counted (Figure 2g). We found that while at E17.5 most, but not all, differentiating HC had no HC neighbors (∼69%), at P0 most differentiating and trans-differentiating HCs had one HC neighbor (∼67% and ∼54% respectively). This result holds also when considering the contact length with neighboring HCs instead of the discrete neighbors count, namely differentiating cells at P0 have longer apical contact length with neighboring HCs compared to E17.5 (Extended Data Figure 3). At both stages differentiating and trans-differentiating HCs had fewer HC neighbors, and smaller contact length with neighboring HCs, compared to the average SC in the tissue (Figure 2g, Extended data Figure 3).

One potential reason for the differences between P0 and E17.5 could arise from the fact that in the more advanced differentiation stage it is less likely to find SCs with no HC neighbors (Figure 2g, ∼22% of all SC have no HC neighbor at E17.5 compared to only ∼5% at P0). To quantitatively assess this effect, we asked what would be the probability of a SC with a given number of HC neighbors to differentiate. To estimate this probability we calculated the percentage of differentiating SCs with zero, one or two HC neighbors, out of every SC with the same number of HC neighbors. The group size of SCs with a given number of HC neighbors was estimated according to the reference mid frame (after 24 hours, same as in Figure 2g). This analysis revealed that a significantly higher fraction of SCs with no HC neighbors differentiate at E17.5 (*∼*19%) compared to P0 (*∼*7%, Figure 2h). The fraction of differentiating SCs with one or two HC neighbors showed no significant difference between E17.5 and P0. Therefore, the observed difference in local differentiation patterns between the two developmental stages stems from the difference in the fraction of differentiating SCs with no HC neighbors.

### Contact-dependent lateral inhibition model captures HCs differentiating next to existing HCs

To better understand what drives the age-dependent differentiation patterns, we computationally investigated possible mechanisms underlying the differences between E17.5 and P0. In particular, while classical lateral inhibition models do not allow HCs differentiating next to existing HCs, Notch signaling has been shown to depend on the size of cell-cell contacts^36^. We therefore performed simulations in which morphological differences between cells were explicitly accounted for, and considered inhibitory signals proportional to the contact length between neighboring cells, leading to heterogeneous inhibitory signals. Relying on a previous contact-dependent lateral inhibition model^36^ we simulate the Notch receptor level, N, the Delta ligand level, D, and a repressor level, R, which is activated by Notch signaling and represses the ligand (Figure 3a). Thus, our model considers a single Notch ligand, Delta (representing both Delta-like1 and Jagged2 from the experimental system^37,38^). To analyze the results of our simulations, we classified high Delta cells and low Delta cells as HCs and SCs, respectively, using the midpoint as a threshold (Supplementary Information section 5.1).

**Figure 3.**
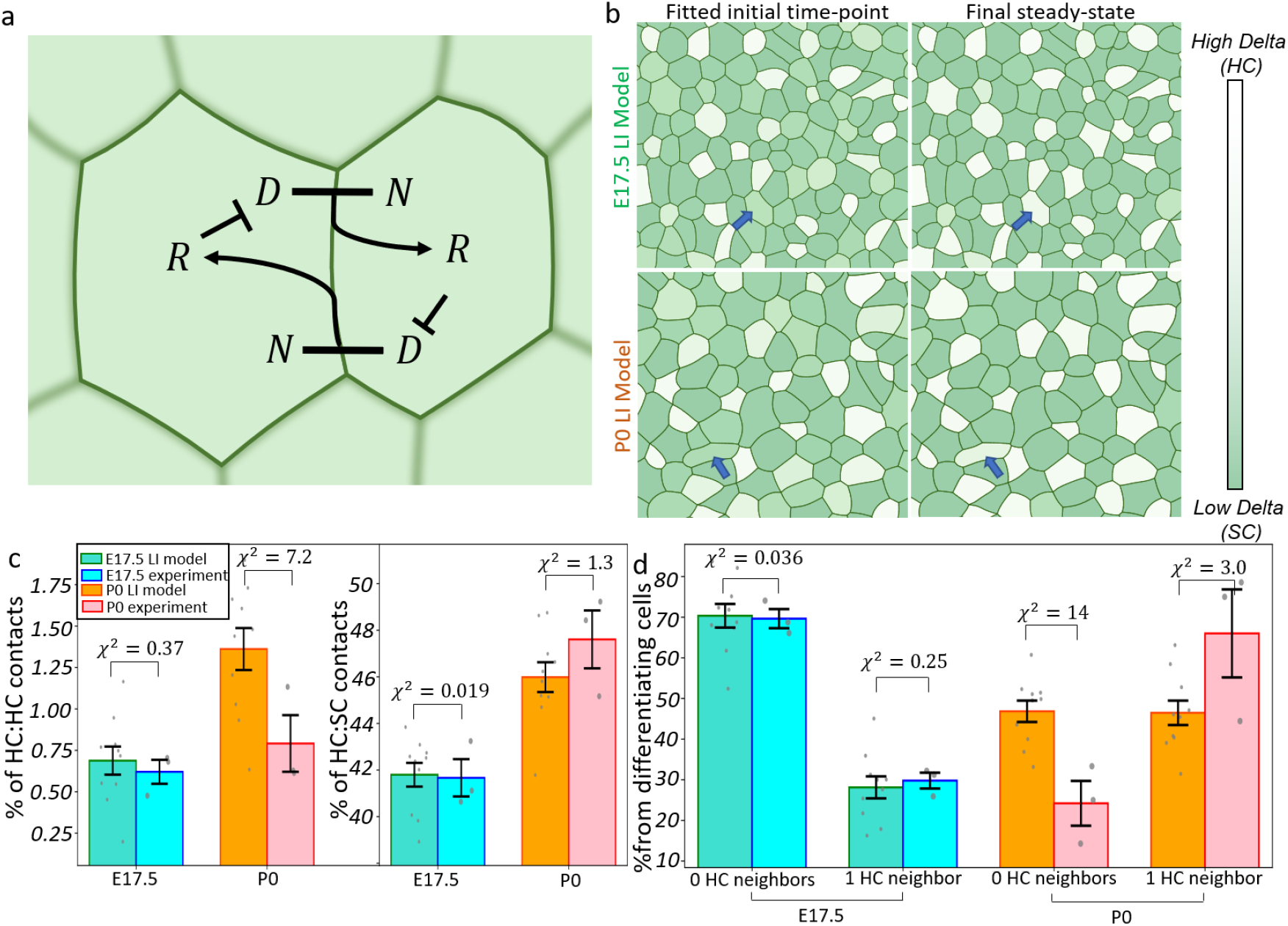
A contact-dependent lateral inhibition model can produce HC differentiation next to existing HCs. **(a)** Model schematics. Interaction of a Notch receptor (N) of one cell with a Notch ligand, Delta (D), in a neighboring cell activates a repressor (R) in the former. This repressor represses the expression of Delta in the same cell, creating a positive feedback loop. In our contact-dependent version, the interaction strength is determined by the length of the contact between the cells. **(b)** Examples of model simulations , with the chosen best matching initial time-points (left) and final steady states (right) for both E17.5 (top) and P0 (bottom) with the best fitting lateral inhibition sensitivity parameters. Both stages produced differentiations next to existing HCs (blue arrows). **(c)** Comparison for the percentage of HC:HC and HC:SC contacts, relative to the total number of contacts, between the first experimental frames and the best matching time point from each model simulation with the best fitting lateral inhibition sensitivity parameters at each stage. **(d)** Comparison for the percentage of differentiating HCs with 0 or 1 HC neighbors between model and experiments, for the best fitting lateral inhibition sensitivity parameters at each stage. The lateral inhibition model is able to produce a good fit to the E17.5 results, but less so for the P0 ones (c-d). Bar heights are determined by group averages, error bars indicate standard errors of the mean (c-d). Scattered points are individual biological repeats for experiments (n=3 for each stage), or initial disordered arrays (n=10 initial arrays, 10 repeats with different initial D,N and R random levels) for model (c-d). The χ2 distance between model and experimental results for each group is indicated (c-d).

To simulate contact-dependent differentiation patterns with data-informed cell morphologies, we quantified the number-of-neighbors distributions from live imaging data, and generated cell arrays with matching statistics (Supplementary Information section 4). For these configurations, we indeed found parameter regimes in which HCs differentiated next to existing HCs (Figure 3b, Extended Data Movie 6-7). A quantitative comparison of the simulations with the experimental data allowed us to identify sensitivity parameters that best fit the experimental results, based on the reduced χ^2^ distance (Figure 3c-d, Methods, Extended Data Table 1). As our total reduced χ^2^ distance is composed of four terms (Figure 3c-d, methods), we consider χ^2^ < 16 (i.e. an average distance less than 2 standard errors) as a good fit. We were able to find fitting parameters that matched the E17.5 patterns (Figure 3c-d left, Extended Data Table 1). We could not, however, find parameters producing a good agreement with the P0 results, even when allowing for stage-specific sensitivity differences (Figure 3c-d right, Extended Data Table 1). Overall, contact-dependent lateral inhibition explains the differentiation of HCs next to existing HCs. However, our results suggest that age-dependent lateral inhibition sensitivity is not sufficient to explain the observed age-dependent differentiation patterns.

### Tissue-scale mechanical properties change between late embryonic and early postnatal stages

Given the evidence that tissue mechanics potentially regulates HC differentiation and regeneration (e.g. the “parking brake hypothesis”^21^), we wondered if changes in tissue mechanics could be the source of the observed differentiation patterns difference between E17.5 and P0. It has been shown that adult mouse utricles have higher mechanical stiffness than embryonic and early postnatal stages^22^, but no significant difference was measured between E17.5 and P0. Similar results have been obtained in the mouse cochlea^39^. However, these measurements did not assess mechanical stress in the tissue. We therefore sought to perform a more sensitive mechanical stress measurement.

To measure the bulk mechanical stress in utricle explants, we used the UV laser ablation system to make a circular cut in the tissue as has been previously performed in other tissues^40^. In all cases, the ablated circle initially shrank, indicating the tissue was mechanically stretched before ablation (Figure 4a,b, Extended Data Movie 8-9). We measured the radius of the shrinking circle over time and found that at E17.5 the cuts shrank faster than at P0, indicating a higher mechanical stress at the former stage (Figure 4c). Treating the tissue as a viscoelastic material, we fitted an exponential function to the changing inner circle radius after ablation for each experiment (Extended Data Figure 4a). We used fit results to calculate the mechanical stress in the tissue^40^, resulting in the bulk stress to viscosity ratio for each stage. We found a significantly higher mechanical stress to viscosity ratio for E17.5 compared to P0 (Figure 4d) indicating a reduction in the mechanical stress, or an increase in viscosity, over time. The initial shrinkage of the ablated circle was followed by a movement of the tissue external to the cut, eventually closing the cut and reforming an intact epithelium within a few minutes (Extended Data Figure 4b). We noticed this contact happened significantly earlier at E17.5 compared to P0 (Extended Data Figure 4b) hinting at a faster healing ability at the early stage.

**Figure 4.**
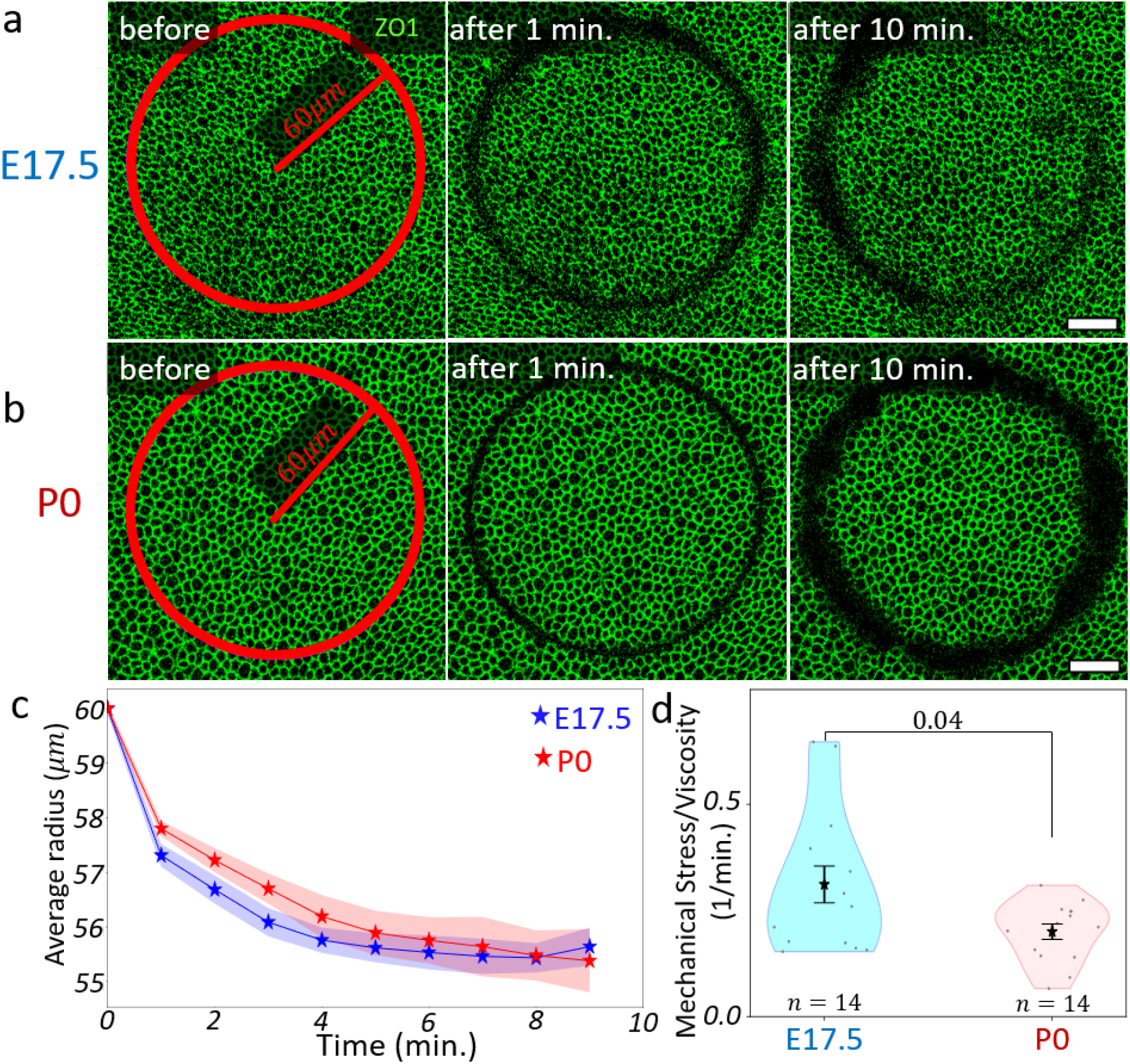
Circular laser ablation reveals higher mechanical stress at late embryonic compared to early postnatal stage. **(a-b)** Film-strips of circular laser ablation experiments for E17.5 (a) and P0 (b) utricle explants. ZO1 (green) marks apical cell boundaries. Circles of 60μm radius were cut in the explants using a high-intensity UV laser and the subsequent dynamics were live-imaged with one minute frame rate. Scale bars - 20μm **(c)** Average inner circle radius over time, averaged over all biological repeats from E17.5 (blue) and P0 (red), indicating an initial faster shrinkage of circular cuts at E17.5 compared with P0 (see Extended Data Figure 4 for individual curves). The shaded area marks standard errors. **(d)** Violin plot of calculated mechanical stress-to-viscosity ratio, indicating a lower mechanical stress, or a higher viscosity, at P0 (red) compared with E17.5 (blue). The number of biological repeats, n, is indicated under each plot. Group averages (stars) and standard errors (error bars) are indicated. p-value was calculated using Welch’s t-test.

### ROCK inhibition reduces local patterning differences between developmental stages

To test whether the observed differences in tissue mechanics can explain the observed HCs differentiation and regeneration patterns, we sought to alter the mechanical stress in the tissue and examine if it had an effect on HC differentiation patterns. To do so, we treated the explants with Y-27632, an inhibitor of Rho-associated coiled-coil protein kinase (ROCK). ROCK is a downstream effector of the Ras homologous (Rho) GTPase signaling pathway which is a known regulator of cytoskeleton arrangement and actomyosin contractility^41^. In the inner ear, it has been shown to be essential for HC and stereocilia development^42,43^.

For this assay, we treated utricle explants with the ROCK inhibitor Y-27632 for 48 hours while performing live imaging and compared the results to untreated control explants (Figure 5, Extended Data Figure 5). We noticed significant morphological differences between treated samples and controls. We observed a significant decrease in HC apical roundness (defined as 4π*⋅area*/*perimeter*^2^) compared to that of SCs (Extended Data Figure 5a-b, Extended Data Movie 10-11). We have previously associated HCs roundness with an increase in mechanical tension on HC cuticular plate^25,26^ (also see hybrid model results in the next section). Thus, the decrease in HC roundness indicates a reduction of mechanical stress in the tissue.

**Figure 5.**
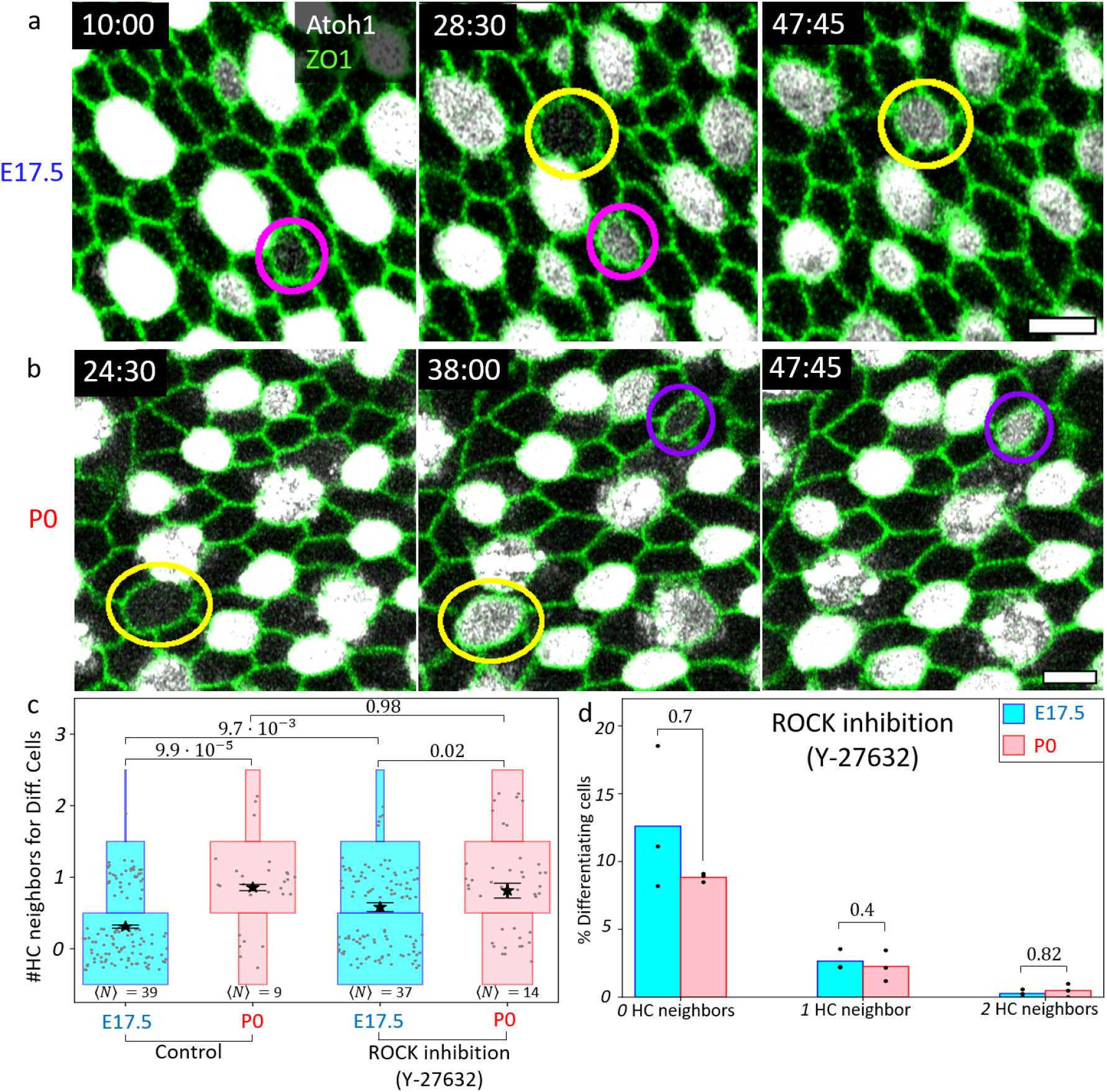
ROCK inhibition reduces patterning differences between late embryonic and early postnatal stages. **(a-b)** Examples of differentiating HCs from live imaging experiments in utricle explants, with ROCK inhibition by 10µM Y-27632 treatment. For both E17.5 (a) and P0 (b) some cells differentiate next to an existing HC, some retain contact with neighboring HC after differentiation (magenta) while others do not (yellow). Some differentiating cells have no HC neighbors (purple). Scale bars - 5µm. **(c)** Discrete violin plot (horizontal histogram) for the number of HC neighbors for each differentiating cell at E17.5 (blue) and P0 (red) for the control (left) and Y-27632 treated (right) explants. While treatment does not induce a significant difference at P0, it does have a significant effect at E17.5: Local differentiation patterns at E17.5 are more similar to those at P0 following Y-27632 treatment. ⟨N⟩, the average total number of cells in each group (over biological repeats), is indicated under each plot. Group means (stars) and standard errors (error bars) are indicated. Scattered data points are randomly shifted along the x- and y-directions. For groups with more than 100 data points only a random subset of 100 points are shown. p-values were calculated by fitting a generalized linear mixed model with Poisson distribution where the biological repeat (3 repeats for each stage at each condition) was treated as the random effect and the group as the fixed effect. **(d)** The percentage of detected differentiations out of the group of SCs with zero, one or two HC neighbors ((# Differentiating cells with n HC neighbors)/(# SCs with n HC neighbors)), for differentiating cells at E17.5 (blue) and P0 (red) for Y-27632 treated explants. Group sizes for each biological repeat were estimated using the data from the reference frame in the middle of each movie. p-values for the difference between E17.5 and P0 are indicated above each pair and were calculated using Mann–Whitney U test.

Next, we analyzed HC differentiation events in both stages. We found that with ROCK inhibition the number of HCs differentiating next to existing HCs significantly increased at E17.5, but did not change at P0 (Figure 5a-c, Extended Data Movie 12-13). Similarly, the contact length of differentiating HCs with their neighboring HCs, increased at E17.5 but not at P0 (Extended Data Figure 5c). Analysis calculating the fraction of differentiating cells out of SCs with 0, 1, or 2 HC neighbors revealed no significant difference between P0 and E17.5 in the treated samples (Figure 5d). Comparing these results to untreated samples (Figure 2h) shows that the main effect of ROCK inhibition is reduction in the differentiation of E17.5 SCs with no HC neighbors. We have attempted to perform single HC ablation experiments under ROCK inhibition. However, we found that the tissue was unable to repair and led to tissue rupture (Extended Data Figure 5d, Extended Data Movie 14). Overall, these data strongly suggest mechanical stress regulates HC differentiation patterns.

### Mechanical stress dependent lateral inhibition model reproduces observed differentiation patterns

Recent works suggested that mechanical properties of the cells can regulate Notch signaling activity^44–46^. To investigate whether this mechanism can explain the age-dependent and mechanically regulated patterns we observe, we performed simulations of an LI model in which the mechanical regulation of Notch signaling is explicitly accounted for. To this end, we combined the contact-dependent^36^ LI model (black symbols in Figure 6a) with a mechanical 2D vertex model^25^, in which cellular morphologies change to minimize the following mechanical energy^47,48^:

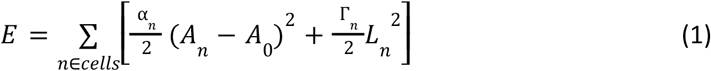

Where *A*_*n*_ , *L*_*n*_ are the apical area and perimeter of the *nth* cell, α_*n*_ is its area elasticity and Γ_*n*_ is its contractility. These morphological changes alter the contact lengths and hence the strength of LI signaling (blue symbols in Figure 6a). We let HCs and SCs have different contractility and area elasticity , which feeds back on cell morphologies^26^ (blue symbols in Figure 6a).

**Figure 6.**
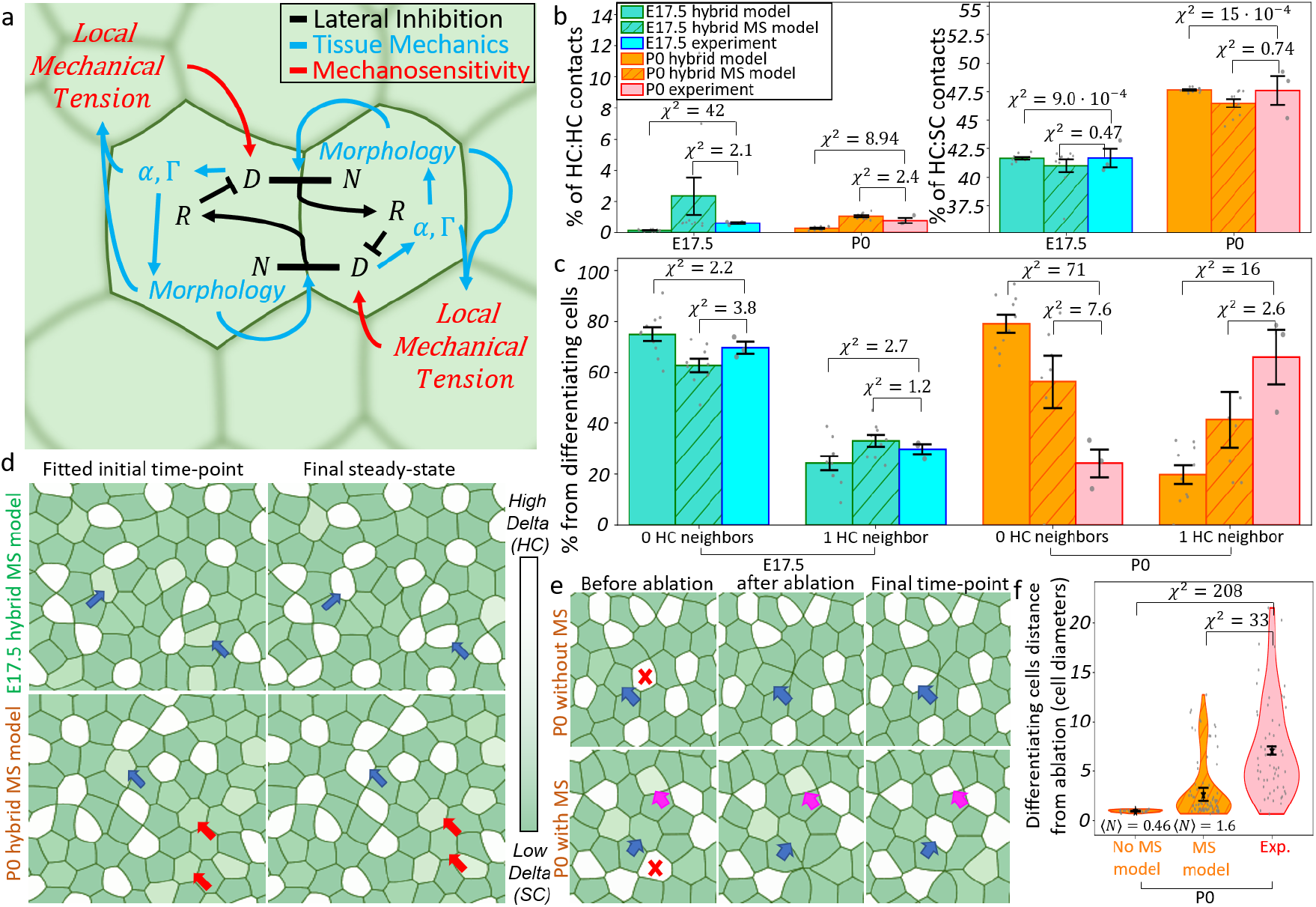
Mechanosensitive lateral inhibition model reproduces observed differentiation patterns. **(a)** Schematics for the model parts: (i) Contact length dependent lateral inhibition (black) with one ligand, Delta (D), one receptor (N) and a repressor (R). (ii) Tissue mechanics (blue) 2D vertex model with changing cell morphology. Differentiation into HC and SC is controlled by Delta and each cell type has different area elasticity (α) and contractility (Γ), affecting mechanical forces and cell morphologies and regulating lateral inhibition (by altering the contact lengths). (iii) Mechanical regulation of Delta production (red) by the local mechanical tension. **(b)** Bar plot comparing the experimental percentage of HC:HC (left) and HC:SC (right) contacts (out of all contacts) with the model results, for each developmental stage, with and without mechanosensitivity. **(c)** Bar plot comparing the experimental number of HC neighbors for each differentiating cell with the model’s results, with and without mechanosensitivity. Bar heights and error bars indicate group averages and their standard errors (b-c). Each scattered point is the result of a single biological repeat for experimental results, or the average of 3 repeats for each initial disordered array for model results (b-c). **(d)** Examples of simulations of the mechanosensitive hybrid model with best fitted parameters. The chosen initial time-points (left) and the final steady states (right) for both E17.5 (top) and P0 (bottom) are presented. With the addition of mechanosensitivity, model results demonstrate many differentiation events next to existing HCs (blue arrows), for both stages. For the P0 simulation, it also shows SCs with no HC neighbors that do not differentiate at steady-state (red arrows). **(e)** Film-strips for single HC ablation simulations with (bottom) and without (top) mechanosensitivity. Without mechanosensitivity, the model predicts a strictly local response where the differentiating cell (blue arrows) is a direct neighbor of the ablated HC (red X). With mechanosensitivity, while some differentiating cells are still the direct neighbors of the ablated HC (blue arrows), others are not (magenta arrows). **(f)** Violin plot comparing the experimental distance between ablated HC and each future differentiating cell, just before ablation, compared with model’s results with and without mechanosensitivity. Each scattered point is a single differentiation event. *⟨*N*⟩* is the average number of differentiations per simulation. Averages (stars) and standard errors (error bars) are shown. Each model was simulated on 10 initial arrays (same for all models, different for each stage) for 5 repeats each (b,c,f). Experimental results include 3 biological repeats for each condition (b, c, f). Reduced χ^2^ distances with experimental distribution for each model are shown (b, c, f). MS - Mechanosensitivity.

To determine the mechanical parameters, we simulated this vertex model first without lateral inhibition dynamics (using fully patterned arrays) and fitted the results to the observed morphologies (Extended Data Figure 6a middle), including cell roundness (HC to SC roundness ratios, Extended Data Figure 6b top), the changes in apical areas in response to laser ablations^25^ (HC to SC area change ratios, Extended Data Figure 6b middle), and the observed shrinkage in the circular ablation experiments (Figure 4c, Extended Data Figure 6b bottom). This fitting process (Supplementary Information section 7) resulted in stage-specific (E17.5 vs P0) values for all mechanical parameters (*A*_0_ for all cells, α_*n*_ , Γ_*n*_ for HCs and SCs, Extended Data Table 2).

To study the effect of direct mechanosensitivity, we incorporated signaling dynamics into the vertex model and introduced a control of Delta production through local membrane tension^49^, such that higher local tension leads to higher Delta production (red symbols at Figure 6a). Assuming stage-independent signaling parameters, the addition of mechanosensitivity introduces a single additional parameter, controlling its strength. We tuned this single parameter by fitting the observed experimental patterns, using the E17.5 sensitivity estimates, for both stages simultaneously (aiming for the lowest sum of the reduced χ^2^ distances for both stages, Extended Data Figure 6a right) and performed a comparative analysis to evaluate the relevance of the direct mechanosensitivity term. We found that the same mechanosensitivity strength could fit the observed patterns for both E17.5 and P0, resulting in an improved agreement with the experimental data compared to the same model without mechanosensitivity (Figure 6b-c, Extended Data Table 1). This improvement was particularly pronounced at P0 (Figure 6b-c, left of each panel, Extended Data Table 1). In addition to capturing the phenomenon of HCs differentiating next to existing HCs, for both stages, without mechanosensitivity (blue arrows in Extended Data Figure 6c, Extended Data Movie 15-16) or with mechanosensitivity (blue arrows in Figure 6d, Extended Data Movie 17-18), including mechanosensitivity also permits capturing the emergence of isolated SCs that do not touch any HCs at P0 (red arrows in Figure 6d, Extended Data Movie 18, quantification in Extended Data Figure 6d). These isolated SCs are still absent for E17.5 (Extended Data Figure 6d). Together, these results suggest that mechanosensitive coupling of lateral inhibition to the mechanical differences between E17.5 and P0 accounts for the observed patterning differences between these developmental stages.

To gain an intuitive understanding of why the addition of direct mechanosensitivity improved model results, we examined the observed cellular morphologies. We found that HCs have consistently rounder apical areas than SCs (Extended Data Figure 5b, Extended Data Figure 6b top), which translated in our model to higher contractility for HCs (Extended Data Table 2). This led to higher tensions at HC:SC interfaces compared to SC:SC contacts (Extended Data Figure 6e). In particular, isolated SCs with no HC neighbors have lower overall tension. The proposed mechanical regulation of Notch signaling then leads to a situation where these isolated SCs have reduced signaling activity and therefore a lower tendency to differentiate into HCs. This effect is more pronounced at P0 since the global mechanical stress is reduced compared to E17.5 (Figure 4d).

Finally, we tested if our model can correctly predict the response to single HC ablation in a parameter-free manner. For that, we simulated the dynamics in response to the ablation of a single randomly chosen HC with and without direct mechanosensitivity. While in the absence of mechanosensitivity, we observed that ablating an HC resulted in the differentiation of one of its direct neighbors (blue arrows in Figure 6e top, Extended Data Movie 19), the addition of mechanosensitivity led to frequent non-neighboring differentiation events (magenta arrows in Figure 6e bottom, Extended Data Movie 20). Quantitative comparison with the experimentally measured distances yielded an improved prediction of the mechanosensitive model, compared to the same model without mechanosensitivity (Figure 6f, Extended Data Table 1). Overall, the mechanosensitive model was able to capture HC differentiation statistics across stages, and to qualitatively predict the response to single HC ablation.

## Discussion

The capacity of HC regeneration in the mouse inner ear is known to decline rapidly in the first few weeks after birth. Here, we used live imaging of vestibular explants to investigate how differentiation dynamics change with age both during normal development and following local HC ablation. The differentiation of HCs and SCs has long been known to be regulated by Notch-mediated LI, where HCs inhibit their neighbors by Notch signaling^10,37,50^. We have observed unexpected differentiation behavior, where new HCs often differentiate next to existing HCs (Figure 2a-d,g). This observation is in contrast to the expected prediction from the classical LI model - a salt-and-pepper pattern where every HC (low Notch activity) is surrounded only by SCs (high Notch activity) ^51,52^. This behavior was age-dependent as we observed a higher fraction of differentiating HCs next to existing HCs at P0 compared to E17.5 (Figure 2g). We further find that the age-dependent difference in differentiation patterns is associated with changes in tissue mechanics between E17.5 and P0 (Figures 4-5). Finally, we show that a hybrid model that takes into account both the regulatory level (i.e. LI) and tissue mechanics (i.e. 2D vertex model) can explain our observations by including contact length-dependency and mechanical regulation of Notch signaling (Figures 3,6).

The emergence of two adjacent low Notch activity cells in the context of LI is rare, but has been reported to occur during the development of sensory organ precursors (SOPs) in Drosophila. In this system, Notch-mediated LI is regulating the differentiation of SOPs from an equivalent group of proneural cells. While in most cases SOPs differentiate more than one cell diameter away^53^, there are a few observations of neighboring SOP pairs in the tissue^53–55^. The emergence of these SOP pairs was attributed to a time delay in the inhibitory signal^56^. Adjacent low Notch cells were also reported in the zebrafish lateral-line^57^, but in this system LI specified a low Notch HC progenitor cell, that later divides into adjacent HCs (for which LI regulates HC polarity)^58,59^. Another system where low Notch cells can form contact is the intestinal stem cell niche, where low Notch Paneth cells form contacts during crypt fission^60^, but these contacts are transient and the cells are already differentiated. Other cases in which adjacent low Notch cells were observed involve Notch signaling perturbations (either genetic or pharmacological)^38,61–63^.

In addition to Notch-mediated LI, we provide evidence that tissue mechanics plays an important role in regulating HC differentiation patterns. In particular, we show a change in the tissue’s mechanical properties between E17.5 and P0 (Figure 4d), and that mechanical stress reduction via ROCK inhibition at E17.5 results in a differentiation pattern resembling that of P0 (Figure 5c-d). These results align with the ‘parking brake hypothesis’, where decreased regeneration capacity in the inner ear was attributed to increased mechanical stiffness^21,22^.

Through a set of simulations parameterized by the experimental observations, we gain quantitative insights into the underlying mechanochemical coupling mechanisms. First, we show that lateral inhibition with contact length dependence^36^ can lead to the differentiation of new HCs next to existing ones, because the disordered tissue morphology provides sufficient signaling variability to allow deviations from strictly alternating patterns^51,52^ (Figure 3). However, morphological coupling alone does not explain all features of the age-dependent differentiation patterns. We found that taking into account the experimentally observed changes in tissue mechanics and introducing an explicit mechanosensitivity of Notch signaling not only fitted the experimental results for both E17.5 and P0, but also accurately predicted the appearance of SCs that do not touch HCs (Extended Data Figure 6d), and the non-local effect of single HC ablation, promoting differentiation of new HCs that are not the direct neighbors of the ablated HC (Figure 6f). Both of these effects are observed experimentally, but not in simulations without mechanosensitivity (Figure 6f, Extended Data Figure 6d). Overall, we find that contact-length dependency and mechanosensitivity of Notch signaling can account for local age-dependent differentiation dynamics observed experimentally.

The assumption that Notch signaling depends on local mechanics is supported by several studies, *in vitro*^44,64,65^ or in organoids^24,66^. In the organoid systems, Notch mechanosensitivity was suggested to be mediated by either ERK^24^ or Hippo^66^ mechanosensitive pathways. In the inner ear, these pathways were linked to HC differentiation and regeneration^24,30,67,68^. In addition, it was shown that nuclear translocation of the Yes-associated protein1 (YAP1), which is a known component of the Hippo pathway^69^, is correlated with regeneration capability in the mammalian vestibular system^70^.

Interestingly, it was recently shown that mechanical regulation of Notch occurs in the developing and regenerating mouse intestine^49^. In this system, local cell density induced the variable expression of YAP1, which controlled Notch-mediated LI. Specifically, they show that YAP1 activates Delta-like1 indirectly through the FOXA1 transcription factor^49^. These findings provide support for indirect Notch mechanosensitivity as a plausible mechanism for HC differentiation in the mammalian utricle. It would be interesting to study whether a similar crosstalk between Hippo and Notch pathways applies in the mammalian utricle as well.

Our experimental approach has some noteworthy limitations. First, our experimental *ex vivo* system might not fully capture the *in vivo* differentiation dynamics, although we did verify that the final patterns do not significantly differ between the two systems (Extended Data Figure 2). Our live imaging assays also focus exclusively on the 2D apical surface, where the alternating HCs and SCs patterns are manifested. In particular, we note that although Notch receptor-ligand interactions usually occur at the apical adherens junctions^71–75^, it could also occur in more basal regions of the tissue^54,76–78^. Finally, to avoid the striolar vs. extrastriolar differences within the utricle^10^, we focused only on the peripheral extra-striolar regions. Future work might focus on differences in regeneration dynamics between these regions.

Our mathematical modeling approach has certain limitations as well. As with the experiments, the model is explicitly 2D, not taking into account the 3D structure of the tissue. Our model considers a simplified version of Notch signaling, including only one ligand and one receptor, without taking into account specific molecular details such as cis-inhibition between receptors and ligands. This simplification is deliberate, as it enables us to draw general conclusions from the model without delving into the complex details.

Our findings might be also relevant to the cochlea, where HC differentiation is controlled by Notch-mediated LI as well^37,38^ and HC regeneration capacity is lower than in the vestibular system^6^. In a recently published work^79^, we have characterized the HC differentiation in response to Notch inhibition and found that only a small sub-population of SCs trans-differentiated at P0-1. This responsiveness was not reflected in the mRNA expression and chromatin accessibility prior to Notch inhibition. Our current findings provide a potential explanation for the reasons underlying this behavior, as the responsiveness might be influenced by the local tissue mechanics.

This work has implications for the pursuit for regenerative treatments for balance and hearing disorders. We suggest that tissue mechanics should be considered as an important factor regulating regeneration. Future treatments might include perturbations of tissue mechanics in a way that would promote HC trans-differentiation. Our study also highlights the general importance of the interplay between local mechanics and signaling in regulating a wide range of developmental and regenerative processes. In particular, such interplay can be driven by the crosstalk between Notch signaling and mechanical forces^45,46,80^ and through the mechanosensitivity of Notch signaling proposed here.

## Methods

### Mice

Rosa26-ZO1-EGFP mice were a gift from Y. Furuta and F. Matsuzaki from RIKEN Laboratory (accession no. CDB0260K) and maintained on a C57BL/6 background. Atoh1-mCherry mice were a gift from C. Petit, Institut Pasteur, and were maintained on a C57BL/6 background. Genotyping was performed using the KAPA HotStart Mouse Genotyping Kit (Sigma-Aldrich, KK7352) using the relevant primers (Extended Data Table 3). For timed mating, one or two females were cross-bred with a male from the second mouse line. The mated females were visually examined to detect pregnancies. E17.5 embryos were dissected 17 days after the day when timed matings were separated. Pregnant females were euthanized using CO2 and embryos were extracted and decapitated into ice-cold phosphate-buffered saline (PBS, Sartorius 02-023-1A). P0 neonates were collected 20 days after timed matings and euthanized by decapitation into ice-cold PBS. As Atoh1-mCherry mice are heterozygous with respect to the insert, embryos and neonates’ brains were examined under a fluorescent microscope (EVOS FL) to detect the mCherry signal. ZO1-EGFP mice are homozygous and therefore all embryos/neonates were positive for the insert. Inner ears were dissected in PBS. Inner ears from Atoh1-mCherry positive mice were used later for live imaging experiments while Atoh1-mCherry negative ears were used for circular ablation. All animal procedures were approved by the Animal Care and Use Committee at Tel Aviv University (approval number TAU-LS-IL-2212-174-2). All institutional guidelines for animal care were strictly followed.

### Utricle Explants Dissection

Utricle explants were dissected in ice-cold PBS using ultra-fine tweezers (Bar Naor BN11252-00). First, protective tissue was extracted to reveal the roof-membrane with its dark cells. Then, the roof-membrane was carefully dissected from the utricle, lateral and anterior cristae. Otoconia were later removed by gently stroking them with the tweezers. Finally, the vestibular nerve was cut and utricles were extracted together with the lateral and anterior cristae and embedded in preincubated (37°C 5% CO2) Matrigel (Corning FAL356237), diluted 15:100 in Dulbecco’s modified Eagle’s medium (DMEM, Sartorius 01-053-1A) on a glass bottom dish. Explants were placed apical side down. Dishes split into four compartments (Hangzhou Xinyou, D35C4-20-1.5-N) were used, to allow for live imaging of up to four explants each time. Embedded explants were then incubated for an additional 5-10 minutes (until Matrigel solidified). 900 µl of explant medium (DMEM supplemented with 1% Fetal Bovine Serum F7524, Biowest S1400-500 1% N-2 supplement, Thermo Fisher 17502048 and 10 µg/ml Ciprofloxacin hydrochloride, Enzo Life Sciences ALX-380-288-G005) was carefully added on top of the embedded explants. Explants were then transferred into a confocal inverted microscope (Zeiss LSM 880) equipped with a life supporting chamber (37°C, 5% CO2) for live imaging. Finally, 100 µl of medium with Y-27632 dihydrochloride (Abcam, ab120129, for ROCK inhibition experiments) or with no additional supplement (for control experiments) were added to the explants, to reach a total medium volume of 1ml. Y-27632 dihydrochloride was diluted to 10mM in PBS and 1µl of the dilution was added to the medium, for a final concentration of 10µM.

### Live Imaging

Explants were live imaged using Zeiss LSM 880 inverted confocal microscope, equipped with an Airyscan detector. We use a 488-nm laser for GFP and 561-nm laser for mCherry. Imaging was performed with a C-Apochromat 40x water-immersion objective with a numerical aperture of 1.2, in Fast Airy-Scan mode. A Z-stack of 40 planes with 0.5 µm separation was acquired for each channel (Atoh1-mCherry and ZO1-EGFP). The striola region for each explant was identified based on its location (in the center of the utricle) and 3D cell morphology (more basal HC nuclei). We avoided these regions and imaged only the extra-striolar peripheral region. A time interval of 15 minutes was used between consecutive frames. This was enough time to acquire the required Z-stacks for up to four explants with a field of view of 200 µm X 200 µm with a resolution of 10 pixels per micron. The equipment was controlled using Zeiss software-”Zen Black”. The resulting movies were processed using Zeiss Airyscan processing.

### Single HC Ablation

To perform HC laser ablation, one frame was acquired first, then 3-4 distant HCs were ablated using Rapp OptoElecronic UGA-42 Caliburn system equipped with a 355-nm UV short-pulse laser. The ablation system was integrated to the microscope’s hardware and software and controlled using Rapp software SysCon. Laser power of 5% and pulse duration of 500 milliseconds was used to ablate the chosen HCs. Ablated HCs were chosen on different field of view corners such that they would be at least 80µ*m* apart. Live imaging was resumed immediately after ablation.

### Circular Ablation

Utricle explants from ZO1-EGFP insert positive mice (without Atoh1-mCherry insert) were used for circular ablation experiments. In these experiments only a single explant was imaged each time, using a regular glass bottom dish (SPL Life Sciences cat no. 200350). Z-stack of 50 planes with one micron separation and a frame rate of one frame per minute was used. To enable the fast frame rate, faster scanning time and lower resolution were used (5.3 pixels per micron). After the first frame was acquired, the mounted laser ablation system was used to ablate a circle with a radius of 60 µm. Laser power of 5% was used with step size 100 and 5 runs. A timer of one minute had been started with ablation and live imaging was resumed exactly when the timer went off.

### Image analysis

Image analysis included a few steps:

1. Apical surface projection-Live imaging experiment results are 3D time-lapse movies. The first analysis step is to detect the apical plane and project the 3D images accordingly, to get 2D time-lapse movies of the apical plane only. As the apical plane is not flat, this surface projection is done by a custom-made algorithm (see code availability section), detecting the apical plane using the ZO1-EGFP signal, which has high intensity at the apical tight-junctions. Briefly, the algorithm constructs a height-map, finding the Z-coordinate of the apical plane for each X,Y location. It starts by detecting a bright spot in the ZO1-EGFP to set the Z for the X,Y starting point, it then expands this height map gradually around this point, keeping the Z difference between adjacent points to be at most one Z-plane.
2. Segmentation- The resulting 2D apical surface movies are segmented based on the projected ZO1-EGFP intensity, using EpySeg^81^. This automatic segmentation is followed by manual corrections, using a custom code in python. Corrected segmentation is then used to find the neighbors for each cell, by looking for cells that share a common apical boundary.
3. Classification-Segmented cells are classified into three options – HC, SC or invalid (areas that are not cells such as holes in the tissue or segmentation errors). This classification is done in three steps - (i) segmented regions that are larger than 100 µ*m*^2^ or smaller than 0.1 µ*m*^2^ are classified as “invalid”. (ii) Segmented regions where more than a given percentage of pixels have Atoh1-mCherry intensity larger than a given percentage from the local maximum intensity are classified as HCs. The area of the local neighborhood to find the local maximum, as well as the threshold intensity and the percentage of required pixels above threshold are chosen manually for each movie. (iii) Manual corrections, done separately for each time-point.
4. Tracking-Tracking is done in two steps - (i) segmented cells centroids are calculated (using region-props) and corrected according to the 2D drift between consecutive time-points (calculated using phase-correlation). (ii) Drift corrected centroids from consecutive frames are linked using TrackPy^82^.
5. Dynamical event detection-Events from the following types are marked manually – division, delamination, differentiation and ablation (for single HC ablation experiments).
6. Data extraction-Relying on segmentation, classification, tracking and event detection, the following data is extracted: (i) The number of dynamical events from each type. (ii) The number of neighbors from each type, and the contact length with each neighbor, for cells undergoing each dynamical event, at the time of the event, and for cells from each type at selected frames. (iii) The area, perimeter and roundness (4π * *area*/*perimeter*^2^) for cells from each type for selected frames.

### Single HC ablation analysis

For single HC ablation experiments, differentiating cells are tracked back to the first frame (before ablation) and the distance to the nearest ablated cell is measured in that frame. For comparison, the distance of every SC to the nearest ablated cell is measured in the same frame. This way, both the differentiating cells distance to ablation and the SCs distance to ablation are dependent on the arbitrary choice of ablated HCs.

### Circular ablation analysis

For each time-point after ablation, the borders of the inner part of the ablated tissue (the inner ablated circle) are fitted into a circle. This is done by manually marking points along this border and fitting them into the circle equation (*x* – *x*_0_)^2^ + (*y* – *y*_0_)^2^ = *R*^2^ via the least-squares method. *x* and *y* are the coordinates of the marked points and *x*_0_ , *y*_0_ , *R* are fit parameters (circle center location and radius respectively). Standard error for the radius is calculated using the standard estimation 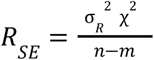 where 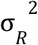 is the fitting error (The diagonal term of the covariance matrix) , χ^2^ is the sum of squared residuals of the least square method, *n* is the number of marked points and *m*=3 is the number of fitting parameters. Next, the change in radius over time is fitted into the following exponential equation *R*(*t*) = (*R*_*start*_ − *R*_*end*_)*e*^−*kt*^ + *R*_*end*_ where *k* (relaxation rate) and *R*_*end*_ (final radius) are fitting parameters. Mechanical stress is calculated based on a previous work^40^ and assuming isotropic stress, by the following equation 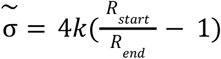, where 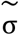 is the mechanical stress normalized by the viscosity. This process is done separately for each experiment to get the distribution of mechanical stress values.

### Visualization and Statistical Analysis

Live imaging results are processed using Fiji^83^. Data visualization is done in python using matplotlib^84^. Statistical analysis is done in python using statsmodels^85^. To avoid confusion, the number of biological repeats is always marked by “n” while the number of data points, in the cases where it is larger than the number of biological repeats (e.g. number of differentiating cells) is marked by “N”. Experiments are classified into one of the following designs - (i) Simple design, where each data point is a biological repeat and only one factor is altered (e.g. comparing circular ablation between developmental stages). (ii) Two by two design, where each data point is a biological repeat but two factors are changing (e.g. comparing the percentage of differentiating HCs with a given number of HC neighbors between developmental stages with or without HC ablation). (iii) Hierarchical design where each data point is a cell within a biological repeat (e.g. comparing number of HC neighbors for each differentiating cell between developmental stages). For each design, statistical tests are performed to choose the best fitting statistical test and calculate p-values. Each design is analyzed by a designated pipeline, as follows:

1. Simple design - For counting data (e.g. number of neighbors) Mann-Whitney U-test is used. For continuous data (e.g. mechanical stress), normality is tested. For large samples (20 < *n*) D’Agostino-Pearson test is used, for medium samples (5 ≤ *n* ≤ 20) Shapiro-Wilk test is used, small samples (*n* < 5) are treated as not-normally distributed. For non-normal distribution of any sample, Mann-Whitney U-test is used. If both samples have normal distribution, Levene’s test is performed to check for equal variances. If variances are equal, Student’s t-test is used. Otherwise, Welch’s-t test is used.
2. Two by two design - Counting data is treated the same as non-normally distributed continuous data. For continuous data, each sample is tested for normality as described for the simple design. For normally distributed data, two-way analysis of variance (ANOVA) with Tukey post-hoc test is used. If any group does not have normal distribution, Scheirer-Ray-Hare test (non-parametric equivalent of two-way ANOVA) is used. If any factor’s effect is statistically significant, but interaction effect is not, Dunn post-hoc test is used. If the interaction effect is significant, pairwise Mann-Whitney U-test with Holm-Bonferroni correction is used.
3. Hierarchical design - If the sample is counting data (e.g. number of neighbors), data is initially fitted using a generalized linear mixed model (GLMM) with Poisson distribution. Then, the fitted model is tested for over-dispersion and zero inflation. If there is over-dispersion and zero inflation, a GLMM with zero-inflated negative binomial distribution is used. If data is zeros inflated, but not over dispersed, GLMM with zero-inflated Poisson distribution is used. If there is only over-dispersion and not zeros-inflation, a GLMM with negative binomial distribution is used. If there is no over-dispersion and no zeros-inflation the Poisson model is retained. For continuous data (e.g. apical contact length) data is tested for normality using Shapiro-Wilk test and for skewness using Fisher-Pearson test. For normal and not skewed data, linear mixed model (LMM) is used. For positive data, Log-transformed LMM is used. For non-normal and skewed data GLMM with inverse Gaussian distribution is used. For non-normal and non-skewed data, GLMM with gamma distribution is used. For all cases, biological repeat is treated as a random effect while checked factor (e.g. developmental stage) is treated as the fixed effect. P-values are calculated for the fixed effect pre-factor difference from zero (i.e. difference from the null-hypothesis which is that the checked factor had no effect).

### Model

#### Lateral inhibition

is modeled based on the following dynamical equations (Supplementary Information section 2):

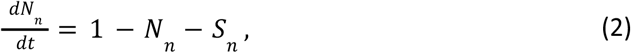

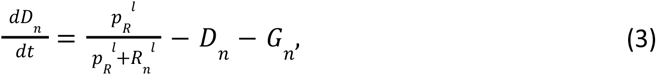

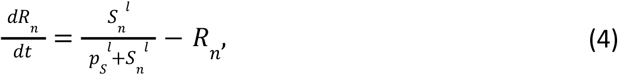

Where *N*_*n*_ , *D*_*n*_ and *R*_*n*_ are the Notch, Delta and repressor concentration in cell *n*, respectively. 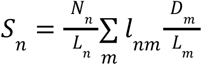 is the total signaling received by cell *n*, 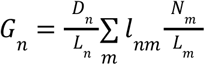 is the total signaling sent by cell *n. L*_*n*_ is cell *n* perimeter, *l*_*nm*_ is the contact length between cells *n* and *m, p*_*R*_ and *p*_*s*_ are the signaling sensitivity parameters, setting the switching point of the Hill-functions, and *l* is the Hill-function exponent, setting the abruptness of the switch. All are non-dimensionalized (Supplementary Information section 2.2-2.4). For simplicity, we set *l* = 3 for all model simulations.

#### Tissue mechanics

is modeled as a 2D vertex model, utilizing the ‘Tyssue’ python package^86^. With a few added functionalities (git reference):

1. Virtual vertices - As performed previously^25,26,87^, virtual vertices (2-way vertices, in addition to 3-way in the original 2D vertex model) are added to allow for more complex cell shapes (Supplementary Figure 1, Supplementary information section 3.2).
2. Periodic boundary conditions - To avoid edge effects, periodic boundary conditions are implemented by stitching the right/upper most edges to the left/lower most edges.
3. Lateral inhibition data - In addition to morphological attributes (e.g. area, perimeter, location), the information about LI related concentrations (Notch, Delta and Repressor) is kept for each cell. This allows for running the 2D vertex model and the LI dynamics in parallel. Including changing mechanical parameters based on the concentrations and changing LI dynamics based on morphology or mechanical forces. Vertex locations are updated according to Eq. 1 and the following equation

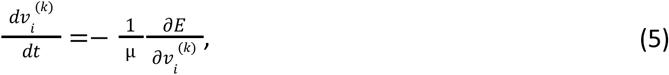

where 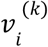 is the *k*′th-coordinate (*k ∈* {*x, y*}) of the *i*′th vertex and µ is the viscosity. In our model, HCs and SCs are attributed with different mechanical parameters. This is done by setting the area elasticity and contractility of each cell (Eq. 1) according to its Delta level:

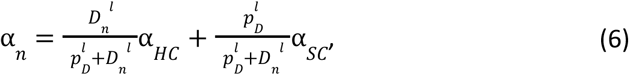

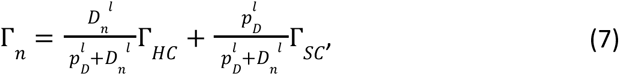

where *p*_*D*_ is the differentiation threshold. α_*HC*_ and α_*SC*_ are the area elasticity for HCs and SCs respectively. Γ_*HC*_ and Γ_*SC*_ are the contractility for HCs and SCs respectively. α_*SC*_ was chosen to be 1 in the non-dimensionalization process (Supplementary Information section 3.4).

#### Direct mechanosensitivity

is implemented by adding mechanical tension activation to equation (3):

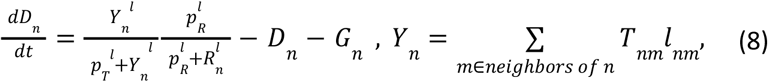

where *T*_*nm*_ is the non-dimensionalized total mechanical tension on the edges between the *n*′th cell and the *m*′th cell and *p*_*T*_ is the mechanosensitivity strength of the model. The mechanical tension for each edge is calculated according to the following equation:

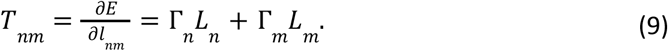

#### Disordered initial configurations

that serve as the initial conditions for the 2D vertex model were created by starting from a hexagonal 20X20 cells arrays and integrating Eq. 1,5 with a randomly chosen mechanical parameters, until reaching a configuration matching the experimental number of neighbors distribution for each cell (Supplementary Figure 2, Supplementary Information section 4)

#### Model simulation

is done in a quasi static fashion, assuming fast mechanical dynamics compared to the LI dynamics. Starting from a chosen initial array, LI levels (D, N and R values for each cell) are randomly initialized for each cell (drawn from a uniform distribution between 0 and 0.01). Then, mechanical parameters are set according to Eqs. 6-7 and vertex positions are updated according to Eqs. 1,5 using forward Euler method with an adaptive dt (Supplementary Information section 3.1), until reaching a steady-state (identified by maximal vertex movement smaller than a given threshold for three consecutive time steps, Supplementary Table 1). Then, the LI levels (D, N and R for each cell) are updated by integrating Eqs. 2-4 for the entire elapsed time (cumulative sum of dt intervals since last LI update). Integration is done using scipy solve_ivp method^88^ with Ronge-Kotta solver (RK45^89,90^). Then, mechanical parameters are updated according to the new Delta levels (Eqs. 6-7) and the quasi static process is repeated. This is done until LI levels reach a steady-state as well (checked in a similar way to mechanical steady-state, Supplementary Table 1). Cell ablation is implemented by manually choosing ablated cells and forcing them to shrink (by changing their *A*_0_ to 0) and delaminated.

#### Fitting model parameters

Generally, our model has seven free parameters: two LI sensitivity parameters - *p*_*S*_ , *p*_*R*_ ; four mechanical parameters – *A*_0_ , α_*HC*_ , Γ_*SC*_ , Γ_*HC*_ ; and a single mechanosensitivity parameter - *p*_*T*_. The differentiation threshold *p*_*D*_ is not fitted, instead it is heuristically chosen by running an LI only model (fixed vertex positions) and calculating the midpoint of the created gap between the high and low delta levels at steady state (Supplementary Figure 3a-b).

Our objective functions are all based on the reduced χ^2^ distance, defined as follows for any observable η:

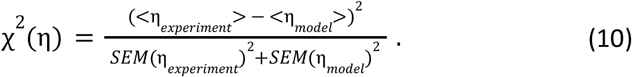

The fitting process for all free parameters is done by adding one layer of complexity at a time, via the following steps (see schematic flow chart in Extended Data Figure 6a):

1. Fitting LI sensitivity parameters - for this step we define the following cost functions:

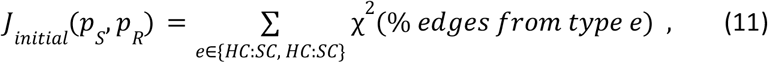

comparing initial patterns, where the experimental values are counted at the first frame of each experiment,

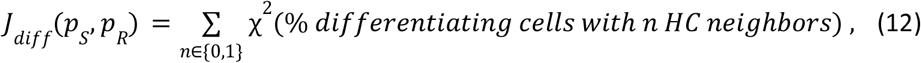

comparing differentiation patterns, and the combined objective function

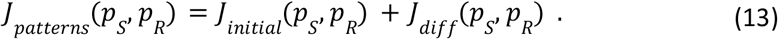

SC:SC edges and differentiating cells with two or more HC neighbors are not accounted for in the objective functions (Eq. 11 and 12 respectively) as they are not independent (as the total percentages add up to 100) and thus carry no extra information. A grid search for *p*_*R*_ and *p*_*S*_ is used to find the (*p*_*R*_ , *p*_*S*_) pair minimizing *J*_*patterns*_ (*p*_*S*_ , *p*_*R*_) (Supplementary Information section 6). The fitting process is done separately for E17.5 and P0 (using the proper initial configurations and experimental results for each stage), to obtain stage specific LI sensitivity parameters. Results show a good fit to the E17.5 results, but not to P0 (Figure 3c-d). Therefore, the best fitting *p*_*S*_ , *p*_*R*_ values for E17.5 were chosen for the hybrid model simulations. These values are fixed from here on, i.e. the same values for *p*_*S*_ , *p*_*R*_ are used for all hybrid model simulations.
2. Fitting mechanical parameters - for this step we use the following objective function

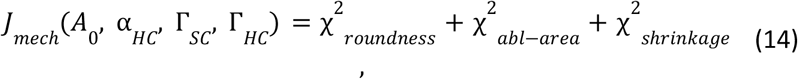

where

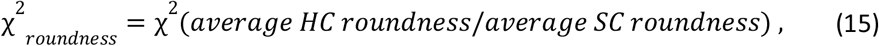

for which the experimental results are obtained from the last frame of every unperturbed live imaging experiment,

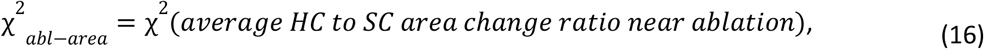

for which the experimental results are obtained from single HC ablation experiments, and

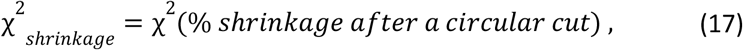

for which the experimental results are obtained from the circular ablation experiments (Figure 4). Shrinkage percentages in the model are estimated by minimizing the energy (Eq. 1) with respect to a uniform linear shrinkage

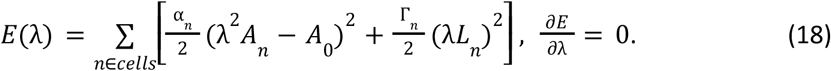 Finding the energy minimizing value of λ for a given steady-state morphology, the shrinkage percentage is given by 100 · (1 − λ). For every combination of *A*_0_ , α_*HC*_ , Γ_*SC*_ and Γ_*HC*_ , the objective function *J*(*A*_0_ , α_*HC*_ , Γ_*SC*_ , Γ_*HC*_) is calculated (Supplementary Information section 7.1-7.4). To search the large parameter space spanned by *A*_0_ , α_*HC*_ , Γ_*SC*_ and Γ_*HC*_ , a heuristic search of the parameter space is done (Supplementary Figure 4, Supplementary Information section 7.5). This search resulted in stage specific mechanical parameter values (Extended Data Table 2).
3. Fitting mechanosensitivity parameter - For this step, the same objective function as for the LI sensitivity parameters are used, but this time as a function of *p*_*T*_ combining the objectives for both developmental stages:

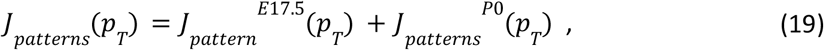

where *J*_*pattern*_ ^*E17*.*5*^ (*p*_*T*_) was calculated via Eq. 11-13 using the experimental E17.5 results and *J*_*pattern*_ ^*PO*^ (*p* _*T*_) was calculated via the same equations, but using the experimental P0 results. To calculate this objective function, a hybrid model was simulated, including both the vertex model and lateral inhibition in parallel (as described in the “model simulation” section). The minimal value for *J*_*pattern*_ (*p*_*T*_) was found using a 1D grid search over different *p*_*T*_ values (Supplementary Information section 8.2-8.3). For each *p*_*T*_ , simulations were performed separately for E17.5 and P0 (using the stage matching initial configurations and fitted mechanical parameters) to calculate *J*_*pattern*_ ^*E17*.*5*^ (*p*_*T*_) and *J*_*patterns*_ ^*po*^ (*p*) respectively (Supplementary Figure 5). The fitted LI sensitivity parameters for E17.5 were used for all simulations (both for E17.5 and P0).

#### Model validation

is done by simulating the hybrid model with the fitted LI sensitivity parameters for E17.5 and the fitted stage specific mechanical parameters. The percentage of SCs with no HC neighbors, out of all cells, was calculated for each resulting steady-state and compared to the experimentally observed percentage for the final frame of each live imaging experiment (Extended Data Figure 6d). Quantitative comparison (Extended Data Table 1) was performed by calculating

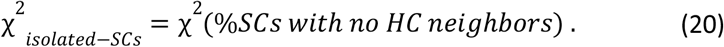

Finally, starting from the resulting steady-state of each P0 simulation without mechanosensitivity, a single HC ablation is simulated for the hybrid model without and with mechanosensitivity, using the P0 fitted mechanical parameters (Supplementary Information section 8.4). For each simulation, one randomly chosen HC is ablated and the subsequent dynamics is simulated, looking for differentiation events. Model results are compared to the experimentally measured distances from the single HC ablation experiments (Main text figure 6f). To translate the experimental results, measured in µ*m*, into cell diameter units, the average experimental cell diameter is estimated using the measured apical areas from the first frames of each single HC ablation (just before ablation, same frame where distances were measured). For this estimation, the circular approximation 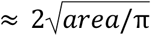 is used. Quantitative comparison (Extended Data Table 1) was performed by calculating

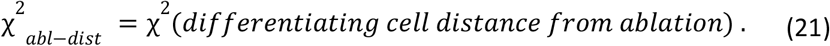

#### Comparing tension on different edges

is done by calculating the simulated mechanical tension for each HC:SC edge and each SC:SC edge at the chosen initial time-point of each simulation using the results of all hybrid model simulations (with and without mechanosensitivity separately). The average tension for each edge type is calculated for each simulation (Extended Data Figure 6e).

## Acknowledgements

We thank the members of the Sprinzak laboratory and K. Avraham for advice and comments on this work. Atoh1-mCherry mice were a kind gift from Christine Petit, Institut Pasteur, France. ZO1-EGFP mice were a kind gift from Yasuhide Furuta and Fumio Matsuzaki from RIKEN, Japan.

## Author contributions

Conceptualization: S.K., M.S., T.D. and D.S. Methodology: S.K., M.S., and D.S. Investigation: S.K., M.S., O.L., T. D., A.E., and D.S. Visualization: S.K., O.L., and D.S. Formal analysis: S.K., M.S., T.D., A.E., and D.S. Supervision: D.S. Writing–original draft: S.K., M.S., T.D., A.E., and D.S. Writing–review and editing: S.K., M.S., T.D., A.E., and D.S. Funding acquisition: D.S.

## Data and materials availability

All data needed to evaluate the conclusions in the paper are present in the paper and/or the Supplementary Materials. Source data and code for image analysis and numerical simulations will be sent upon request. Requests for the Atoh1-mCherry mice should be submitted to. Requests for the ZO1-EGFP mice should be submitted to. These can be provided pending scientific review and a completed material transfer agreement.

