## Extended data for "Age-dependent mechanical stress underlies unexpected vestibular differentiation and regeneration patterns"

### Extended Data Figures

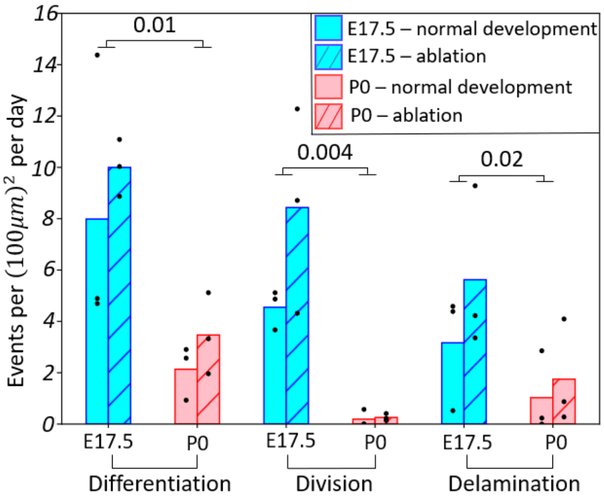


**Extended Data Figure 1. Frequency of differentiation, division, and delamination events.** The number of events from each type (Differentiation into HC, cell division, and delamination) per $(100\mu m)^2$ per day at each live imaging experiment for normal development (uniform fill) and after HC ablation (dashed lines fill) at E17.5 (blue) and P0 (red). Bars show group averages, each point represents a single biological repeat (n=3 for each condition). P-values were calculated using Scheirer-Ray-Hare test (non-parametric two-way ANOVA) to compare developmental stages. Interaction effect was not statistically significant in any case.

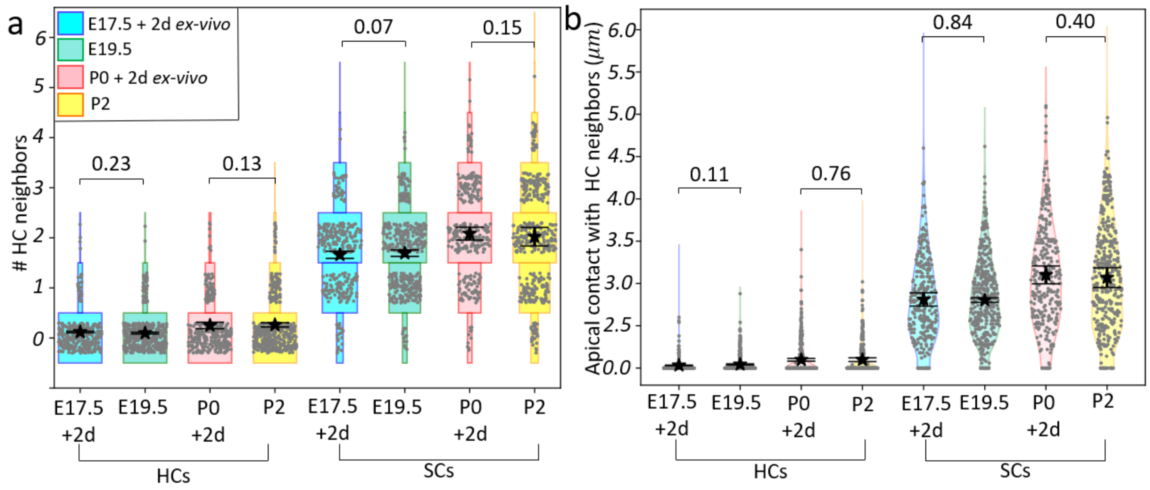


**Extended Data Figure 2. Similar differentiation patterns are observed *ex vivo* and *in vivo*. (a)** A discrete violin plot (horizontal histogram) for the number of HC neighbors for HCs (left) and SCs (right). No significant statistical differences were detected between *ex vivo* utricles (last movie frame, after two days of live imaging) and *in vivo* ones (taken immediately after dissection at the matching developmental stage). **(b)** A violin plot for the apical contact lengths with HC neighbors for HCs (left) and SCs (right). No significant statistical differences were detected between *ex vivo* and *in vivo* utricles. Group averages (stars) and standard errors (error bars) are indicated. P-values were calculated by fitting a generalized linear mixed model (Methods). N>900 cells for all groups. The number of biological repeats is n=3 and n=4 for each *ex vivo* and *in vivo* developmental stage, respectively.

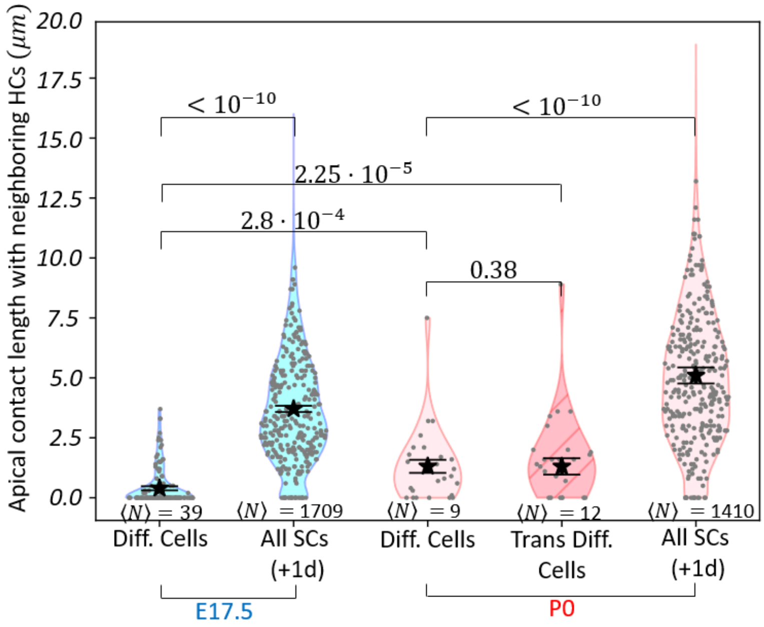


**Extended Data Figure 3. Differentiating and trans-differentiating HCs contact length with neighboring HCs.** Violin plot of the total contact length of each cell with its neighboring HCs for differentiating cells (E17.5, blue and P0, red) and trans-differentiating cells (P0 only, dashed red), compared with all SCs at a reference frame (chosen at the middle of each movie, after 24 hours of live imaging). Group average (stars) and SE (error bars) are indicated. ⟨N⟩, the average number of cells in each group (over biological repeats) is indicated under each plot. p-values were calculated by fitting a generalized linear mixed model with Gamma or Inverse Gaussian distribution (depending on data skewness) where the biological repeat (three for each condition) was treated as the random effect and the group label (e.g. differentiating cells vs. all SC) as the fixed effect.

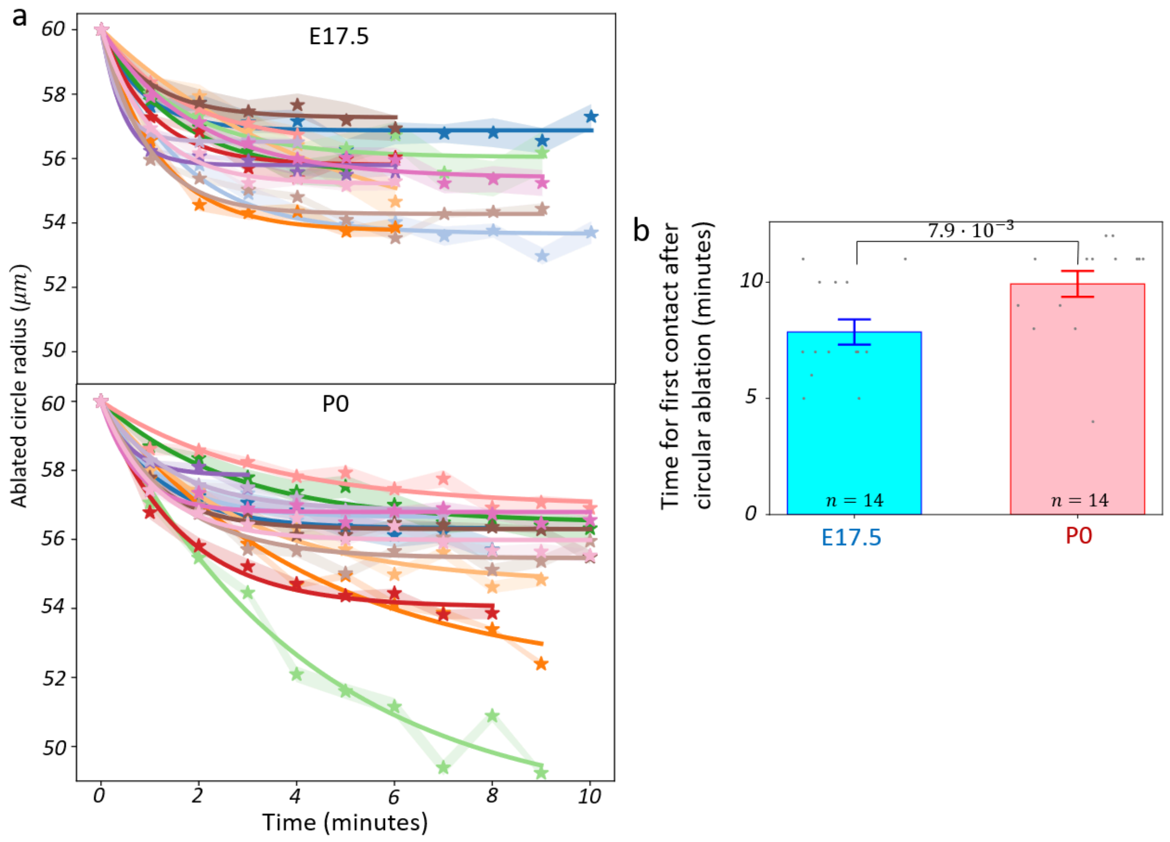


**Extended Data Figure 4. Circular ablation supplementary data.(a)** Exponential fits (solid lines) for the ablated circle radius shrinkage over time (stars) at E17.5 (top) and P0 (bottom). Each color represents a separate biological repeat. Radii were calculated by fitting the ablated shape to a circle for each time point. Shaded areas indicate the estimated standard errors for the radii. **(b)** Comparison of the time from circular ablation to the first contact of the ablated circle with the external part of the tissue indicates faster closing for the E17.5 utricles compared with P0. The number of biological repeats, n, is indicated under each bar. Bar heights are group averages and error bars indicate standard errors. P-value was calculated using Mann–Whitney U test.

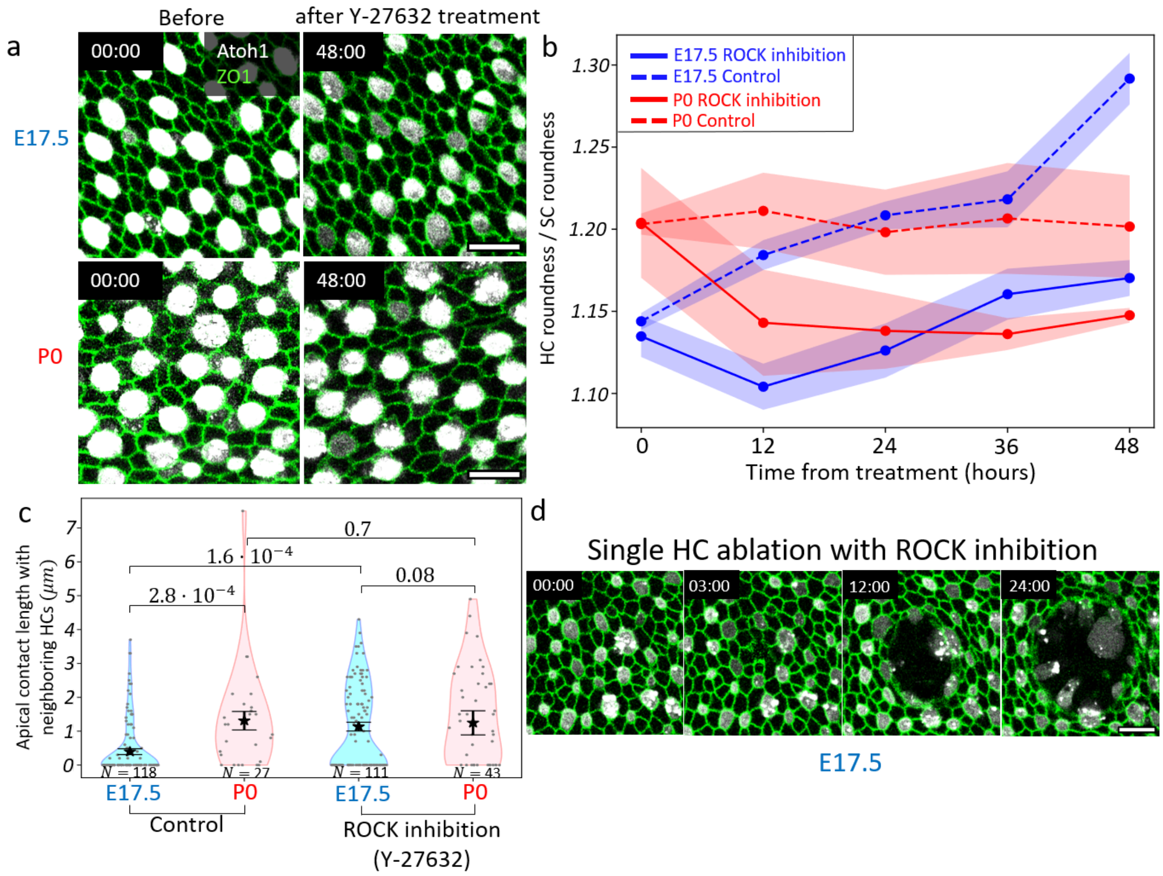


**Extended Data Figure 5. ROCK inhibition supplementary data. (a)** Live imaging examples of utricle explants with ROCK inhibition by 10μM Y-27632 treatment. 48 hours post-treatment the shapes of HCs and SCs are more similar to each other (right panels) than they were before treatment (left panels), when HCs appeared to be more round. Scale bars - 10μm. **(b)** Average HC roundness ($4\pi \cdot Area/Perimeter^2$) divided by average SC roundness as a function of time after Y-27632 treatment (solid lines) or without a treatment (dashed lines) for E17.5 (blue) and P0 (red) utricle explants. ROCK inhibition results in a lower HC roundness to SC roundness ratio than in the control samples. Each data point is the average of three biological repeats (with the exception of the 48 hours timepoint for control samples which includes only two biological repeats for E17.5). Shaded area marks the standard errors. **(c)** Violin plot for the apical contact length with HC neighbors for each differentiating cell at E17.5 (blue) and P0 (red) for the control (left) and Y-27632 treated (right) explants. While treatment does not induce a significant difference at P0, it does have a significant effect for E17.5 local patterns, which have no significant difference compared to P0 patterns after treatment. ⟨N⟩, The average total number of cells in each group (over biological repeats), is indicated under each plot. Group means (stars) and standard errors (error bars) are indicated. Scattered data points are randomly shifted along the x-directions. For groups with more than 100 data points only a random subset of 100 points are shown. p-values were calculated by fitting a generalized linear mixed model with Inverse Gaussian or Gamma distribution (chosen based on sample skewness) where biological repeats (three for each stage at each condition) were treated as the random effect and the group as the fixed effect. **(d)** Live imaging of a single HC ablation after ROCK inhibition. The explant was treated with Y-27632 just before ablation. Unlike untreated explants (Figure 2c) the tissue does not heal after ablation. Instead, a hole in the tissue is created and grows with time. Time after ablation is indicated. Scale bar - 10 μm.

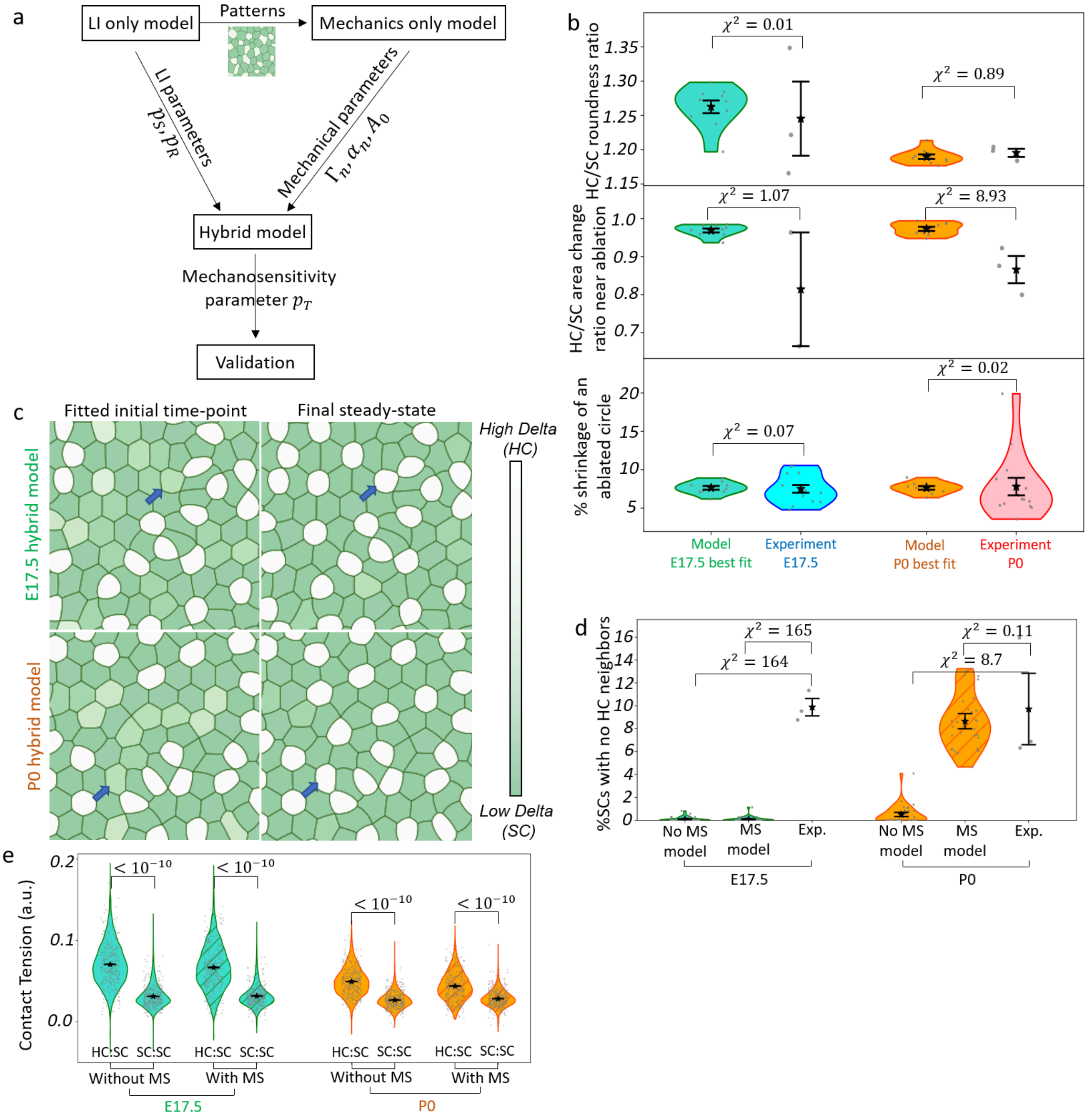


**Extended Data Figure 6. Model results supplementary data. (a)** Schematic flow chart for the fitting process used to find model parameters. Each box shows what version of the model was simulated, what was its input (in-going arrows), and what was the output of this stage after comparing its results with the experimental data (out-going arrows). **(b)** Results of the mechanical parameters fitting procedure for the three fitted morphological characteristics – The ratio of HC to SC average roundness at the final time-point (top), the ratio of the change in HC to SC area for cells that are a direct neighbor of an ablated cell (middle), and the percentage of shrinkage after a circular laser ablation (bottom). Each scattered point is the result of an individual simulation, for the model results (one for each of the 10 initial arrays), or a biological repeat for the experimental results (n=14 for shrinkage. n=3 for roundness and ablation, except for ablation at E17.5 where n=2. . **(c)** Simulations of the hybrid model without mechanosensitivity. The chosen initial time-points (left) and final steady states (right) for both E17.5 (top) and P0 (bottom) are presented. Both stages produced differentiations next to existing HCs (blue arrows). **(d)** Violin plot comparing the experimental percentage of isolated SCs (% SCs with no HC neighbors out of all cells) at the final experimental frames to model simulations. The model simulations show the percentage of isolated SCs in the final steady states, with and without mechanosensitivity. Model results show a negligible amount of these isolated SCs with E17.5 fitted parameters, even with mechanosensitivity (left). For P0 fitted parameters, some isolated SCs are produced. More isolated SCs are produced with mechanosensitivity, matching the experimental results. **(e)** Violin plot for the distributions of simulated mechanical tension, comparing the tension for HC:SC and SC:SC contacts with and without mechanosensitivity. Overall, the simulated tension is significantly larger for the HC:SC contacts compared to the SC:SC contacts. The p-values for the compared pairs are shown and calculated by fitting a linear mixed model with Log-Transformed distribution. Scattered points are the results of individual simulations (3 repeats for each of the 10 initial arrays for each model) or a single biological repeat (n=3 each stage) for experiments (b, d). For groups with fewer than 10 data points, only averages, standard errors and scattered points are displayed (b, d). Averages (stars) and standard errors (error bars) are shown (b, d, e). Reduced $\chi^2$ distances with the relevant experimental result is shown above each model result (b, d). MS – Mechanosensitivity.

#### Extended Data Movies Captions

**Extended Data Movie 1. Live imaging dynamical events example.** E17.5 utricle explant exhibiting one cell division event (red), two delamination events (blue) and one differentiation event (magenta). ZO1-EGFP (apical boundaries) is shown in green. Atoh1-mCherry (hair cells) is shown in white. Live imaging time is indicated. Scale bar - 5μm.

**Extended Data Movie 2. HC differentiation example from an E17.5 utricle.** Live imaging of differentiating HC (magenta). Live imaging time is indicated. Scale bar - 5μm.

**Extended Data Movie 3. HC differentiation example from a P0 utricle.** Live imaging of differentiating HC (magenta). Live imaging time is indicated. Scale bar - 5μm.

**Extended Data Movie 4. HC differentiation after single HC ablation example from an E17.5 utricle.** Live imaging of HC ablation (red X) followed by a nearby HC differentiation (magenta). Time from ablation is indicated. Scale bar - 5μm**.**

**Extended Data Movie 5. HC differentiation after single HC ablation example from an P0 utricle.** Live imaging of HC ablation (red X) followed by a nearby HC differentiation (magenta). Time from ablation is indicated. Scale bar - 5μm**.**

**Extended Data Movie 6. Lateral inhibition model example E17.5.** An example for the contact-dependent lateral inhibition model simulation results with E17.5 fitted initial array and the best fitting lateral inhibition parameters. Cells are colored based on their Delta levels. Low Delta cells (SCs) are green while high Delta cells (HCs) are white.

**Extended Data Movie 7. Lateral inhibition model example P0.** An example for the contact-dependent lateral inhibition model simulation results with P0 fitted initial array and the best fitting lateral inhibition parameters. Cells are colored based on their Delta levels. Low Delta cells (SCs) are green while high Delta cells (HCs) are white.

**Extended Data Movie 8. Circular ablation example from an E17.5 utricle.** Live imaging of circular ablation at E17.5. Time from ablation is indicated. Ablated circle (red) radius - 60μm. ZO1-EGFP is shown in green. Scale bar - 20μm.

**Extended Data Movie 9. Circular ablation example from a P0 utricle .** Live imaging of circular ablation at P0. Time from ablation is indicated. Ablated circle (red) radius - 60μm. ZO1-EGFP is shown in green. Scale bar - 20μm.

**Extended Data Movie 10. ROCK inhibition example from an E17.5 utricle.** Live imaging of E17.5 utricle explant after Y-27632 treatment. Time from treatment is indicated. Scale bar - 10μm.

**Extended Data Movie 11. ROCK inhibition example from a P0 utricle.** Live imaging of P0 utricle explant after Y-27632 treatment. Time from treatment is indicated. Scale bar - 10µm.

**Extended Data Movie 12. HC differentiation after ROCK inhibition example from an E17.5 utricle.** Live imaging of two differentiating cells for a Y-27632 treated explant at E17.5. One differentiating cell retains contact with its neighboring HC (magenta) while the second one does not (yellow). Time from treatment is indicated. Scale bar - 5µm.

**Extended Data Movie 13. HC differentiation after ROCK inhibition example for a P0 utricle.** Live imaging of two differentiating cells for a Y-27632 treated explant at P0. One cell differentiates next to another HC and later loses contact (yellow) while the second one differentiates with no HC neighbors (purple). Time from treatment is indicated. Scale bar - 5µm.

**Extended Data Movie 14. Single HC ablation combined with ROCK inhibition example from an E17.5 utricle.** Live imaging of a single HC ablation at E17.5, performed immediately after Y-27632 treatment. Time from ablation is indicated. Scale bar - 10µm.
STAR methods

**Extended Data Movie 15. hybrid model example E17.5.** An example for the hybrid model, without mechanosensitivity, simulation results with E17.5 fitted initial array, the best fitting E17.5 lateral inhibition parameters from the Lateral inhibition only model and the best fitting E17.5 mechanical parameters. Cells are colored based on their Delta levels. Low Delta cells (SCs) are green while high Delta cells (HCs) are white.

**Extended Data Movie 16. hybrid model example P0.** An example for the hybrid model, without mechanosensitivity, simulation results with P0 fitted initial array, the best fitting E17.5 lateral inhibition parameters from the lateral inhibition only model and the best fitting P0 mechanical parameters. Cells are colored based on their Delta levels. Low Delta cells (SCs) are green while high Delta cells (HCs) are white.

**Extended Data Movie 17. hybrid mechanosensitive model example E17.5.** An example for the hybrid model with mechanosensitivity, simulation results with E17.5 fitted initial array, the best fitting E17.5 lateral inhibition parameters from the lateral inhibition only model, the best fitting E17.5 mechanical parameters and the best fitting mechanosensitivity parameter for both stages. Cells are colored based on their Delta levels. Low Delta cells (SCs) are green while high Delta cells (HCs) are white.

**Extended Data Movie 18. hybrid mechanosensitive model example P0.** An example for the hybrid model with mechanosensitivity, simulation results with P0 fitted initial array, the best fitting E17.5 lateral inhibition parameters from the lateral inhibition only model, the best fitting P0 mechanical parameters and the best fitting mechanosensitivity parameter for both stages. Cells are colored based on their Delta levels. Low Delta cells (SCs) are green while high Delta cells (HCs) are white.

**Extended Data Movie 19. Single HC ablation hybrid model example.** An example for single HC ablation simulation results for the hybrid model, without mechanosensitivity. Ablation (red X) resulted in a differentiation of an SC that was a direct neighbor of the ablated HC. Simulated with the best fitting E17.5 lateral inhibition parameters from the lateral inhibition only model and the best fitting P0 mechanical parameters, starting from a steady state produced by the same model version (with the same parameters). Cells are colored based on their Delta levels. Low Delta cells (SCs) are green while high Delta cells (HCs) are white.

**Extended Data Movie 20. Single HC ablation hybrid mechanosensitive example.** An example for single HC ablation simulation results for the hybrid model, with mechanosensitivity. Ablation (red X) resulted in a differentiation of two SCs, one was a direct neighbor of the ablated HC while the other one was not. Simulated with the best fitting E17.5 lateral inhibition parameters from the lateral inhibition only model, the best fitting P0 mechanical parameters and the best fitting mechanosensitivity parameter for both stages, starting from a steady state produced by the same model version (with the same parameters). Cells are colored based on their Delta levels. Low Delta cells (SCs) are green while high Delta cells (HCs) are white.

#### Extended Data Tables

| Model version | Stage | $p_R$ | $p_S$ | $p_T$ | $J_{initial}$ | $J_{diff}$ | $J_{patterns}$ | $\chi^2_{isolatedSCs}$ | $\chi^2_{abl-dist}$ |
|---|---|---|---|---|---|---|---|---|---|
| LI only | E17.5 | 0.3 | 0.1 | - | 0.386 | 0.285 | 0.671 | 163.4 | - |
| LI only | P0 | 0.25 | 0.15 | - | 8.525 | 16.765 | 25.290 | 9.66 | - |
| Hybrid | E17.5 | 0.3 | 0.1 | 0 | 42.189 | 4.871 | 47.060 | 164.4 | - |
| Hybrid | P0 | 0.3 | 0.1 | 0 | 8.943 | 87.405 | 96.348 | 8.70 | 207.6 |
| Hybrid with MS | E17.5 | 0.3 | 0.1 | 0.08556 | 2.545 | 5.011 | 7.557 | 164.97 | - |
| Hybrid with MS | P0 | 0.3 | 0.1 | 0.08556 | 3.111 | 10.156 | 13.267 | 0.913 | 33.51 |

**Extended Data Table 1. Model scores summary.** Model reduced $\chi^2$ scores compared with the relevant experimental results for each model version with its best fitting parameters. MS - Mechanosensitivity.

| Stage | $\Gamma_{SC}$ | $\Gamma_{HC}$ | $\alpha_{HC}$ | $A_0$ | $\chi^2_{roundness}$ | $\chi^2_{abl-area}$ | $\chi^2_{shrinkage}$ | $J_{mech}$ |
|---|---|---|---|---|---|---|---|---|
| E17.5 | 0.0105 | 0.070 | 3.5 | 0.74181 | 0.0102 | 1.068 | 0.072 | 1.242 |
| P0 | 0.0105 | 0.043 | 1.757 | 0.75854 | 0.888 | 8.927 | 0.023 | 9.837 |

**Extended Data Table 2. Mechanical fit summary.** Mechanical fit reduced $\chi^2$ scores and best fitting parameters for each stage.

| **Name** | **Forward** | **Reverse** |
|---|---|---|
| Rosa26-ZO1-WT 370 | TATCAGTAAGGGAGCTGCAGT | ACCCCAGATGACTACCTATCCT |
| Rosa26-ZO1-WT 448 | TCCCAAAGTCGCTCTGAGTT | GAAAATACTCCGAGGCGGATCA |
| Rosa26-ZO1-EGFP-insert | CAAGGTGAACTTCAAGATCCGC | ACCCCAGATGACTACCTATCCT |
| Atoh1Cherry-XFPN | GGTAGTTTGCCGTAATGTGAGTG | CTCCTCGCCCTTGCTCACC |
| Atoh1Cherry-U-L | CGAGGAAGCGCAGCCTTCCAG | CGGCGGGGTGCTTCACGTAG |

**Extended Data Table 3. Mouse genotyping primers.** Listing the forward and reverse primers used to genotype the transgenic mouse lines.
