## Supplementary Information for "Age-dependent mechanical stress underlies unexpected vestibular differentiation and regeneration patterns"

### Supplementary Information – Theory and computational analysis for a contact-length dependent and mechanically regulated lateral inhibition model

#### 1. Model overview: Notch signaling model links lateral inhibition parameters to mechanically regulated patterning

To investigate how lateral inhibition and tissue mechanics together regulate pattern formation in the developing utricle, we developed a theoretical framework that couples Notch signaling dynamics to cell mechanical properties and tissue morphology (Main Text Figure 6a). The goal of the model is to test whether developmental changes in cell mechanical properties can alter HC patterning while the core lateral inhibition parameters remain fixed.

We developed the model progressively. We first considered a contact length dependent lateral inhibition model on fixed irregular morphologies. It extends the classical lateral inhibition framework by allowing the signaling strength between neighboring cells to depend on their shared contact length. This reduced model provides a constrained setting for identifying lateral inhibition parameters that reproduce the experimentally observed HC neighbor statistics.

We then extended the model to include changing tissue morphology using a 2D vertex model. In this framework, differentiation into hair cells (HCs) or supporting cells (SCs) changes the mechanical properties of cells, including their contractility and area elasticity . These changes modify cell shapes, contact lengths, and tissue topology, which in turn feed back onto signaling through the contact length dependent interactions. We find the mechanical parameters best fitting each stage by comparing the model's resulting morphologies to experimental results (Extended Data Figure 6b).

Finally, we incorporated a direct coupling between mechanical stress and signaling, motivated by the experimentally observed stress dependent behavior. Specifically, we added positive regulation of Delta production by local mechanical stress. In this mechanosensitive model, mechanics can influence patterning in two ways: indirectly, by changing cell-cell contact geometry, and directly, by modulating Delta production (Main Text Figure 6a).

At each stage, we asked whether the model can reproduce key experimental features of HC patterning, including HCs that differentiate adjacent to existing HCs and SCs that do not neighbor any HCs (Main Text Figure 2g-h). We then tested whether changing mechanical parameters, while keeping the lateral inhibition parameters fixed, can shift the model between patterning regimes resembling different developmental stages. Finally, we examined whether the mechanically coupled model can reproduce the single HC laser ablation results (Main Text Figure 2e).

#### 2. Contact-length dependent lateral inhibition model

##### 2.1 Contact length scales Notch-Delta signaling between neighboring cells

The first extension to the classical lateral inhibition framework introduces contact length dependent signaling. Notch-ligand interactions occur along cell-cell interfaces, suggesting that the strength of signaling between neighboring cells depends on the length of their shared contact, as shown in our previous work[1]. We therefore asked whether incorporating this geometric effect alone could account for the deviations from classical salt-and-pepper patterns observed in the utricle.

We modeled contact length dependent signaling using the following equations:

$$\frac{dn_{nm}}{dt} = \frac{\beta_N}{L_n} - \gamma_N n_{nm} - \frac{\kappa^+ \kappa^s}{\kappa^- + \kappa^s} n_{nm} d_{mn}, \tag{1}$$

$$\frac{dd_{nm}}{dt} = \frac{\beta_D}{L_n} \frac{p_R^l}{p_R^l + R_n^l} - \gamma_D d_{nm} - \frac{\kappa^+ \kappa^s}{\kappa^- + \kappa^s} n_{mn} d_{nm}, \tag{2}$$

$$\frac{dR_n}{dt} = \beta_R \frac{\left( \sum_{m \in neighbors\ of\ n} \frac{1}{\gamma_S} \frac{\kappa^+ \kappa^s}{\kappa^- + \kappa^s} n_{nm} d_{mn} l_{nm} \right)^l}{p_S^l + \left( \sum_{m \in neighbors\ of\ n} \frac{1}{\gamma_S} \frac{\kappa^+ \kappa^s}{\kappa^- + \kappa^s} n_{nm} d_{mn} l_{nm} \right)^l} - \gamma_R R_n. \tag{3}$$

Here $n_{nm}$, $d_{nm}$ are Notch and Delta concentrations of cell $n$ along the edge with cell $m$. $R_n$ represents the repressor level of cell $n$. $L_n$ denotes the perimeter of cell $n$, and $l_{nm}$ is the contact length between cells $n$ and $m$. The parameters $\beta_N$, $\beta_D$, $\beta_R$ are the production rates of Notch, Delta and Repressor, while $\gamma_N$, $\gamma_D$, $\gamma_R$ are their degradation rates. The parameters $\kappa^+$, $\kappa^-$ describe the association and dissociation constants of Notch and Delta respectively. $\kappa^S$ is the rate associated with conversion of the Notch-Delta complex into a signal (namely, the inverse time it takes for Notch to be cleaved, once it interacts with Delta, releasing NICD). $\gamma_S$ is the degradation rate of the signal (i.e. the cleaved NICD). The exponent $l$ is the Hill coefficient, and $p_R$, $p_S$ are Hill threshold parameters setting the location of the steepest response.

#### 2.2 Non-dimensionalization reduces the number of independent parameters

To simplify interpretation and reduce the number of independent parameters, we non-dimensionalized the equations:

$$\frac{d\tilde{n}_{nm}}{d\tilde{t}} = \frac{1}{\tilde{L}_n} - \tau_N \tilde{n}_{nm} - \tilde{n}_{nm} \tilde{d}_{mn}, \tag{4}$$

$$\frac{d\tilde{d}_{nm}}{d\tilde{t}} = \frac{1}{\tilde{L}_n} \frac{\tilde{p}_R^l}{\tilde{p}_R^l + \tilde{R}_n^l} - \tau_D \tilde{d}_{nm} - \beta \tilde{n}_{mn} \tilde{d}_{nm}, \tag{5}$$

$$\frac{d\tilde{R}_n}{d\tilde{t}} = \frac{\left( \sum_{m \in neighbors\ of\ n} \tilde{n}_{nm} \tilde{d}_{mn} \tilde{l}_{nm} \right)^l}{\tilde{p}_S^l + \left( \sum_{m \in neighbors\ of\ n} \tilde{n}_{nm} \tilde{d}_{mn} \tilde{l}_{nm} \right)^l} - \tilde{R}_n, \tag{6}$$

where the dimensionless variables were defined as follows: $\tilde{\beta} = \frac{\beta_N}{\beta_D}$, $\kappa = \frac{\kappa^+\kappa^s}{\kappa^- + \kappa^s}$, $L_0 = \frac{\kappa\beta_D}{\gamma_R^2}$, $\tilde{t} = \gamma_R t$, $\tilde{n}_{nm} = \frac{\kappa}{\gamma_R\tilde{\beta}} n_{nm}$, $\tilde{d}_{nm} = \frac{\kappa}{\gamma_R} d_{nm}$, $\tilde{R}_n = \frac{\gamma_R}{\beta_R} R_n$, $\tau_N = \frac{\gamma_N}{\gamma_R}$, $\tau_D = \frac{\gamma_D}{\gamma_R}$, $\tilde{L}_n = \frac{L_n}{L_0}$, $\tilde{l}_{nm} = \frac{l_{nm}}{L_0}$, $\tilde{p}_S = \frac{\gamma_S}{\beta_N} p_S$, $\tilde{p}_R = \frac{\gamma_R}{\beta_R} p_R$.

##### 2.3 Uniform membrane distribution reduces the edge-based model to cell-level equations

To further simplify the model, we assumed that Notch and Delta are uniformly distributed along the cell apical boundaries. This assumption reflects that these proteins are produced inside the cell and subsequently transported to the membrane, where they are distributed across the entire perimeter rather than localized to specific interfaces. We further assumed this distribution happens faster than all other relevant time scales in the system (e.g. binding, degradation etc.). Under this assumption, the edge specific concentrations $n_{nm}$ and $d_{nm}$ could be expressed in terms of the total Notch and Delta levels in each cell, significantly reducing the number of equations in the system. We therefore defined the total levels of Notch and Delta in cell $n$ as (using the dimensionless variables):

$$\tilde{N}_n \equiv \sum_{m \in neighbors\ of\ n} \tilde{n}_{nm}\tilde{l}_{nm}\ ;\ \tilde{D}_n \equiv \sum_{m \in neighbors\ of\ n} \tilde{d}_{nm}\tilde{l}_{nm}.$$

We could also define the signal and the trans-signal in cell $n$ as

$$\tilde{S}_n \equiv \sum_{m \in neighbors\ of\ n} \tilde{l}_{nm}\tilde{n}_{nm}\tilde{d}_{mn}\ ;\ \tilde{G}_n \equiv \sum_{m \in neighbors\ of\ n} \tilde{l}_{nm}\tilde{n}_{mn}\tilde{d}_{nm}.$$

Under the assumption of uniform membrane distribution, the edge-specific concentrations satisfy

$$\tilde{n}_{nm} = \frac{\tilde{N}_n}{\tilde{L}_n}\ ;\ \tilde{d}_{nm} = \frac{\tilde{D}_n}{\tilde{L}_n}.$$

So that the signal terms could be written as

$$\tilde{S}_n = \frac{\tilde{N}_n}{\tilde{L}_n} \sum_{m \in neighbors\ of\ n} \tilde{l}_{nm}\frac{\tilde{D}_m}{\tilde{L}_m}\ ;\ \tilde{G}_n = \frac{\tilde{D}_n}{\tilde{L}_n} \sum_{m \in neighbors\ of\ n} \tilde{l}_{nm}\frac{\tilde{N}_m}{\tilde{L}_m}.$$

Substituting these expressions into Eqs. (4–6) yields the reduced system:

$$\frac{d\tilde{N}_n}{d\tilde{t}} = \sum_{m \in neighbors\ of\ n} \tilde{l}_{nm}\frac{d\tilde{n}_{nm}}{dt} = 1 - \tau_N\tilde{N}_n - \frac{\bar{L}}{l_s}\tilde{S}_n, \quad (7)$$

$$\frac{d\tilde{D}_n}{d\tilde{t}} = \sum_{m \in neighbors\ of\ n} \tilde{l}_{nm}\frac{d\tilde{d}_{nm}}{dt} = \frac{\tilde{p}_R}{\tilde{p}_R + \tilde{R}_n} - \tau_D\tilde{D}_n - \tilde{\beta}\tilde{G}_n, \quad (8)$$

$$\frac{d\tilde{R}_n}{d\tilde{t}} = \frac{\tilde{S}_n^{\,l}}{\tilde{p}_S + \tilde{S}_n^{\,l}} - \tilde{R}_n. \quad (9)$$

This reduction decreased the size of the system from $2N_{edges} + N_{cells}$ equations to $3N_{cells}$ equations.

##### 2.4 Parameter choices for the lateral inhibition model

To reduce the number of independent parameters, we set the length scale $L_0$ to be the average cell perimeter in the initial morphology. We also set $\tau_N = \tau_D = 1$, corresponding to equal degradation rates for Notch, Delta and the repressor, and $\tilde{\beta} = 1$, corresponding to equal production rates of Notch and Delta. We set $l = 3$ for computational convenience. This leaves two undetermined parameters in the lateral inhibition model $\tilde{p}_S$ and $\tilde{p}_R$, representing the sensitivity of Repressor and Delta production to the activation or inhibition signals, respectively.

#### 3. Simulating tissue mechanics using 2D vertex model

As our experimental results indicate that there are mechanical differences between E17.5 and P0, and that these differences may regulate differentiation, resulting in the age dependent differentiation patterns observed. We therefore sought to incorporate tissue mechanics in our model. To do that we developed a 2D vertex model, based on the existing "tyssue" python package[2]. In the existing package (and in 2D vertex models in general), cells in the tissue are modeled as 2D polygons. The polygon vertex locations are the dynamical variables of the model. They are updated according to the following equation

$$\frac{dv_i^{(k)}}{dt} = -\frac{1}{\mu}\frac{\partial E}{\partial v_i^{(k)}}, \qquad (10)$$

where $v_i^{(k)}$ is the $k$'th coordinate ($k \in \{x, y\}$) of the $i$'th vertex. $\mu$ is the viscosity and $E$ is the mechanical energy of the tissue. For the energy of the tissue, we chose the commonly used form

$$E = \sum_{n \in cells}\left[\frac{\alpha_n}{2}(A_n - A_{0n})^2 + \frac{\Gamma_n}{2}L_n^{\ 2}\right]. \qquad (11)$$

The first term is area elasticity. $A_n$ is the $n$'th cell area, calculated by vertex locations. $A_{0n}$ is its preferred area, a fixed parameter. $\alpha_n$ is its area elasticity, determining how easy it would be for it to deviate from its preferred area. The second term is perimeter contractility. $L_n$ is the $n$'th cell perimeter, calculated by vertex locations. $\Gamma_n$ is its contractility, determining how much it would like to minimize its perimeter.

##### 3.1 Adaptive dt

To simulate tissue mechanics we used a forward Euler method. Tissue morphology is updated according to Eq. 10-11, with an adaptive time interval dt. Simulation starts with an initial dt. After each simulation step, the longest distance a single vertex has moved is calculated. If this distance is larger than a given threshold (Supplementary Table 1), or if an edge crossing was detected, this step is reversed and dt is decreased by a given factor (Supplementary Table 1). Time interval dt can decrease down to a given threshold, below it the simulation would stop and be declared as a failure (Supplementary Table 1). After a successful step, dt would increase by a given factor (Supplementary Table 1) until reaching its initial value.

##### 3.2 Virtual vertices addition to allow for non polygonal morphologies

The shape of cells' apical surface in our tissues is not necessarily a polygon with straight edges, and in some cases not even convex (Supplementary Figure 1a). In order to allow for these shapes in our model, we split straight polygonal edges by the addition of mid edge vertices, which we termed "virtual vertices" (Supplementary Figure 1b). The addition and removal of virtual vertices is done by setting a maximal and minimal bond lengths. If an edge length increases above the maximal bond length, it is evenly split by a virtual vertex addition. If an edge length decreases below the minimal bond length, it is merged with a neighboring edge by virtual vertex removal (if this edge connects two virtual vertices, one of them is chosen for removal at random).

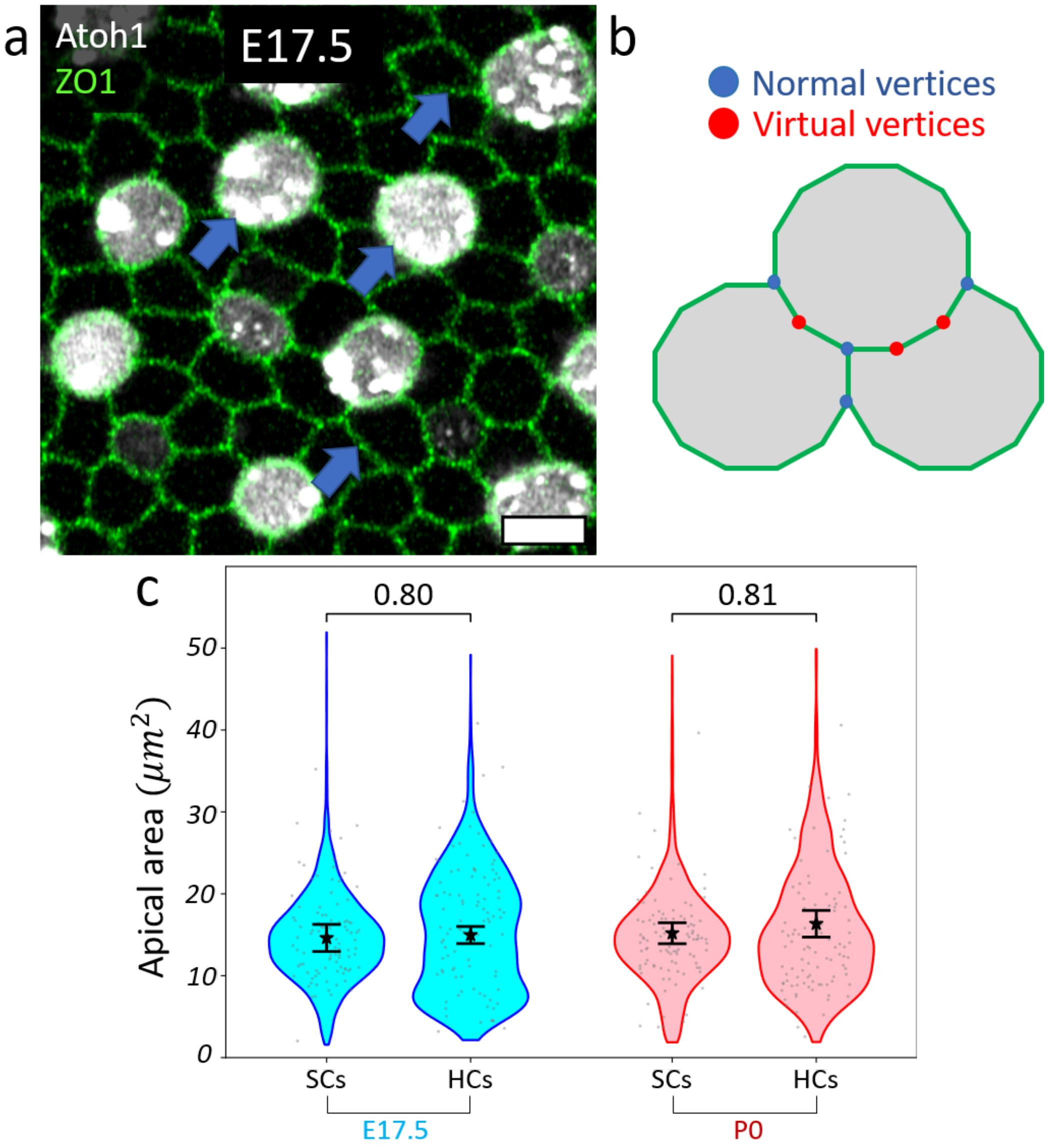


**Supplementary Figure 1. Model design supplementary information. (a)** Experimental example from E17.5 utricle explant's apical plane showing non straight cell boundaries (blue arrows). Atoh1 (HC marker) is shown in white and ZO1 (apical boundaries) in green. Scale bar - 5µm **(b)** Adding virtual vertices (red) in addition to the normal tricellular junction vertices (blue) allows for rounder and non-convex cell shapes. **(c)** a violin plot comparing the measured apical area of HCs and SCs, at the final frame of each live-imaging experiment (normal development) for each stage. Apical areas of HCs and SCs show no significant difference for both stages. Group averages are marked by starts. Error bars show standard errors of the mean. Each scattered point is the apical area of an individual cell, only 100 points are shown for each group. Group sizes - N>300 for each group at each biological repeat. p-values were calculated using a LMM with n=3 repeats for each condition. HCs – hair cells. SCs - supporting cells.

##### 3.3 Topological events

Our model accounts for four kinds of topological events:

1. Intercalations - short edges (below intercalation threshold, Supplementary Table 1) undergo intercalation (T1 transition[3]).
2. Cell divisions - cells with area above division threshold (Supplementary Table 1) divide at a random angle.
3. Delaminations - cells with area below delamination threshold (Supplementary Table 1) are removed from the array.
4. Ablation - cells chosen manually for ablation are forced to shrink (and subsequently delaminate) by setting their $A_0$ to 0.

To avoid the occasional creation of very sharp angles, vertices with sharp angles (below the sharp angle threshold, Supplementary Table 1) are removed either by virtual vertex removal (if possible) or by intercalation.

##### 3.4 Simplifying assumptions and non-dimensionalization

As the mean apical area of HCs and SCs does not differ significantly (Supplementary Figure 1c) we made preferred area uniform by setting $A_{0n} = A_0$. The remaining parameters, $\alpha_n$ and $\Gamma_n$, are set according to cell type. In our case we consider two cell types, HCs and SCs. Cell types are set based on Delta concentration with a smooth transition, using the following equations:

$$\alpha_n = \frac{D_n^l}{p_D^l + D_n^l}\alpha_{HC} + \frac{p_D^l}{p_D^l + D_n^l}\alpha_{SC}, \tag{12}$$

$$\Gamma_n = \frac{D_n^l}{p_D^l + D_n^l}\Gamma_{HC} + \frac{p_D^l}{p_D^l + D_n^l}\Gamma_{SC}, \tag{13}$$

where $p_D$ is the differentiation threshold, i.e. the critical Delta level that separates high Delta HCs from low Delta SCs (see section 5.1 below discussing the determination of $p_D$ value). $\alpha_{SC}$ was chosen to be 1 in the non-dimensionalization process.

To work with dimensionless variables, we defined $\tilde{\alpha}_{HC} = \frac{\alpha_{HC}}{\alpha_{SC}}, \tilde{\Gamma}_{HC} = \frac{\Gamma_{HC}}{\alpha_{SC}a^2}, \tilde{\Gamma}_{SC} = \frac{\Gamma_{SC}}{\alpha_{SC}a^2}$, $\tilde{\tilde{A}}_0 = \frac{A_0}{a^2}, \tilde{\tilde{A}}_n = \frac{A_n}{a^2}, \tilde{\tilde{L}}_n = \frac{L_n}{a}, \tilde{E} = \frac{E}{\alpha_{SC}a^4}, \tilde{\mu} = \frac{\gamma_R}{\alpha_{SC}a^2}\mu, \tilde{v}_i^{(k)} = \frac{v_i^{(k)}}{a}$.

Where $a$ is the mean initial distance between neighboring cells' centroids (the "lattice constant") which sets the length-scale for the 2D vertex model. Note that lengths while length in the 2D vertex model are measured in units of $a$, in the lateral inhibition model they are measured in units of $L_0$ (section 2.2). Therefore, when passing contact lengths, perimeters and areas from the vertex model to the lateral inhibition model, we converted them via multiplication by $\frac{a}{L_0}$. We used the double-tilde signs in $\tilde{\tilde{A}}_0$, $\tilde{\tilde{A}}_n$ and $\tilde{\tilde{L}}_n$ to mark this difference in normalization. With these definitions we get the non-dimensional equations:

$$\frac{d\tilde{v}_i^{(k)}}{d\tilde{t}} = -\frac{1}{\tilde{\mu}}\frac{\partial \tilde{E}}{\partial \tilde{v}_i^{(k)}}, \tag{14}$$

$$\tilde{E} = \sum_{n \in cells}\left[\frac{\tilde{\alpha}_n}{2}(\tilde{\tilde{A}}_n - \tilde{\tilde{A}}_0)^2 + \frac{\tilde{\Gamma}_n}{2}\tilde{\tilde{L}}_n^2\right]. \tag{15}$$

The type dependent parameters are now set by the following equations

$$\tilde{\alpha}_n = \frac{\tilde{D}_n^l}{\tilde{p}_D + \tilde{D}_n^l}\tilde{\alpha}_{HC} + \frac{\tilde{p}_D}{\tilde{p}_D + \tilde{D}_n^l}, \tag{16}$$

$$\tilde{\Gamma}_n = \frac{\tilde{D}_n^l}{\tilde{p}_D + \tilde{D}_n^l}\tilde{\Gamma}_{HC} + \frac{\tilde{p}_D}{\tilde{p}_D + \tilde{D}_n^l}\tilde{\Gamma}_{SC}. \tag{17}$$

To conclude, the mechanical 2D vertex model has four mechanical parameters: $\tilde{\tilde{A}}_0$, $\tilde{\alpha}_{HC}$, $\tilde{\Gamma}_{HC}$ and $\tilde{\Gamma}_{SC}$ which can either have fixed values (e.g. mechanical parameters fit, section 7) or dynamical values updated according to the Delta levels (e.g. hybrid model, section 8).

**3.5 Mechanosensitivity**

Notch direct mechanosensitivity was implemented by adding a direct influence of tissue mechanics on lateral inhibition. Specifically, we hypothesized a mechanosensitive activator of Delta, $Y$, which is directly controlled by the membrane tension. Assuming a fast response of this activator to the changing mechanical tension, such that Y levels are always in steady-state with respect to the current mechanical tensions:

$$Y_n = \sum_{m \in neighbors\ of\ n} T_{nm} l_{nm}, \tag{18}$$

where $l_{nm}$ is the apical contact length between cells $n$ and $m$ and $T_{nm}$ is the tension on that boundary. We added this to the lateral inhibition model by updating Eq. 8 (with the $\tau_D = \tilde{\beta} = 1$ simplifying assumptions) to the following form:

$$\frac{d\tilde{D}_n}{d\tilde{t}} = \frac{Y_n^{\,l}}{p_T^{\,l} + Y_n^{\,l}} \frac{\tilde{p}_R^{\,J}}{\tilde{p}_R^{\,J} + \tilde{R}_n^{\,J}} - \tilde{D}_n - \tilde{G}_n, \tag{19}$$

where $p_T$ is mechanosensitivity threshold.

##### 3.6 Calculating mechanical tension

The mechanical tension on the edge between cell m and cell n is given by

$$T_{nm} = \frac{\partial E}{\partial l_{nm}} = \Gamma_n L_n + \Gamma_m L_m\ . \tag{20}$$

where the partial derivatives of the perimeter is given by

$$\frac{\partial L_k}{\partial l_{nm}} = \frac{\partial}{\partial l_{nm}} \sum_{j \in cell\ k\ neighbors} l_{kj}\ , \tag{21}$$

which is 1 if k=m or k=n and cell $m$ is a neighbor of cell $n$ and 0 otherwise.

##### 3.7 Non-dimensionalization of mechanosensitivity equations

For non-dimensionalization, we defined the non-dimensional parameters $\tilde{T}_{nm} = \frac{T_{nm}}{\alpha_{SC} a^3}$, $\tilde{p}_T = \frac{p_T}{\alpha_{SC} a^3 L_0}$, $\tilde{Y}_n = \frac{Y_n}{\alpha_{SC} a^3 L_0}$ which gives the non-dimensional equations

$$\tilde{T}_{nm} = \tilde{\Gamma}_n \tilde{\tilde{L}}_n + \tilde{\Gamma}_m \tilde{\tilde{L}}_m, \tag{22}$$

$$\frac{d\tilde{D}_n}{d\tilde{t}} = \frac{\tilde{Y}_n^{\,l}}{\tilde{p}_T^{\,l} + \tilde{Y}_n^{\,l}} \frac{\tilde{p}_R^{\,J}}{\tilde{p}_R^{\,J} + \tilde{R}_n^{\,J}} - \tilde{D}_n - \tilde{G}_n\,;\ \tilde{Y}_n = \sum_{m \in neighbors\ of\ n} \tilde{T}_{nm} \tilde{l}_{nm}. \tag{23}$$

#### 4. Generation and validation of developmental-stage-specific configurations

To generate the initial tissue configurations, later used for all simulations, we produced synthetic irregular morphologies independently of the Notch-Delta dynamics. The morphologies were generated using the 2D vertex model described in section 3, with each simulation initialized from a regular hexagonal lattice. The lattice constant, defined as the initial distance between neighboring cell centroids, was set to $a = 1$. To progressively disrupt the initial hexagonal organization, each cell $n$ was assigned independently sampled random values of $\tilde{\tilde{A}}_{0n}$(drawn from a uniform distribution between 0 and 1) and $\tilde{\Gamma}_n$ (drawn from a uniform distribution between 0 and 0.2). $\tilde{\alpha}_n$ was set to its SC value 1. The model was then evolved for one dimensionless time unit, after which new cell specific values of these parameters were sampled. This was done for 25 time units. This repeated parameter resampling, mechanical relaxation, and topological transitions generated disordered, non-hexagonal arrays (Supplementary Figure 2a).

These randomly sampled, cell specific mechanical parameters were used only to generate diverse irregular morphologies. They were not retained in the subsequent simulations of the model, in which the mechanical parameters were assigned according to the assumptions described in section 3.

At each saved time point, we compared the simulated distribution of cell neighbor numbers with the corresponding experimental distributions measured at E17.5 and P0 (Supplementary Figure 2b). For each developmental stage, the time point yielding the highest p-value in the statistical comparison, calculated by fitting a generalized linear mixed model (GLMM) with Poisson distribution to the data, was selected as the best matching morphology. (Supplementary Figure 2c, Supplementary Table 2). If the best matching morphology had been achieved at the end of the simulation, the simulation was resumed for 5 additional time units, to make room for further improvement. This procedure was repeated independently to generate a set of 10 developmental stage specific morphologies for E17.5 and P0. From here on, cell divisions were disabled in all models.

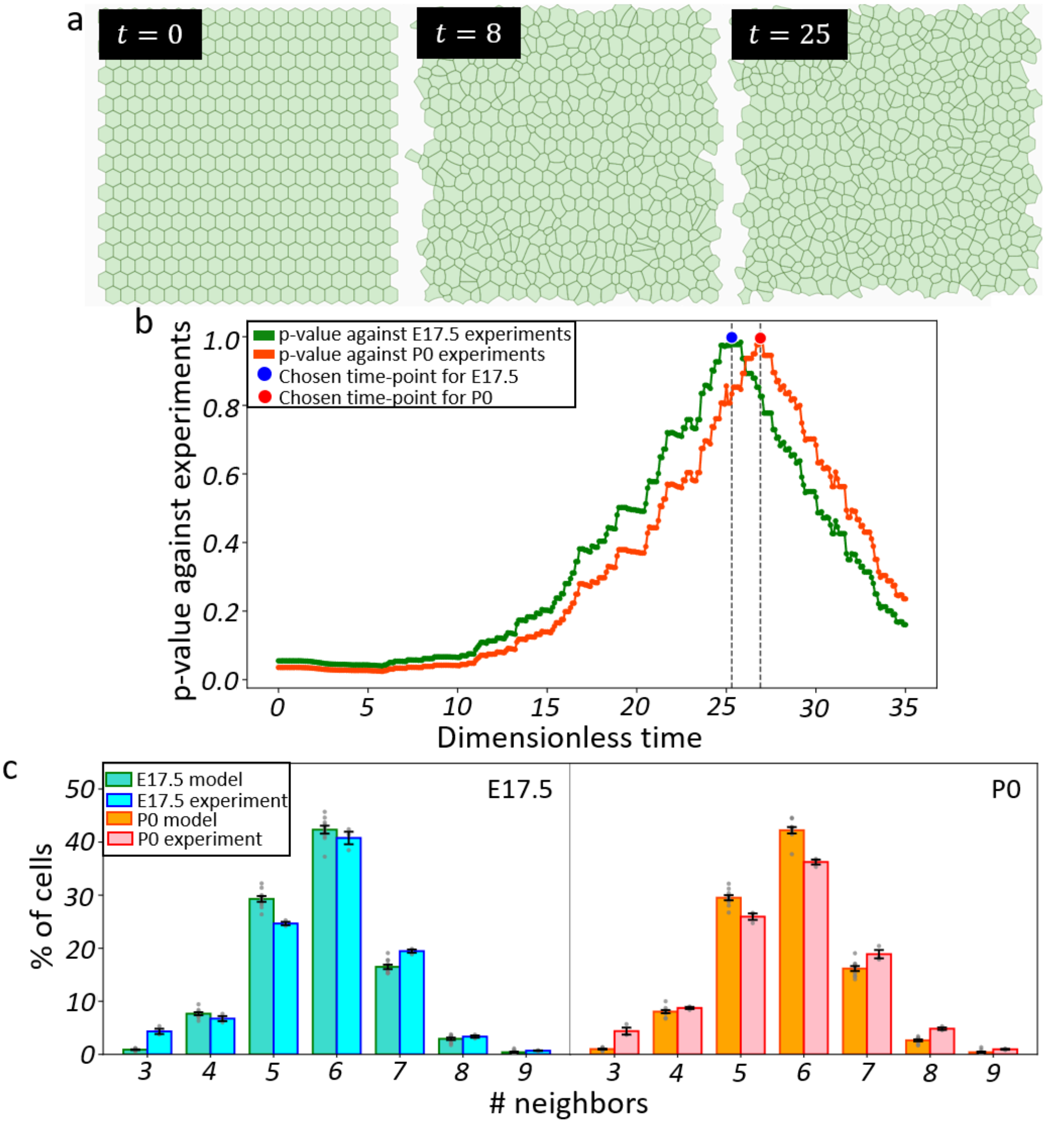


**Supplementary Figure 2. Generating developmental stage dependent initial disordered configurations. (a)** Disordered configuration is generated by starting from a hexagonal array (left) and simulating 2D vertex mechanics with randomly changing mechanical parameters, and cell divisions, to achieve progressively disordered configurations (middle and right). **(b)** Each time-point is compared with the experiments by calculating the p-value between the model's number of neighbors distribution for each cell, and the experimental one, at the first frame of each live imaging experiment. This comparison is done separately for E17.5 (green) and P0 (orange) experimental results. The time point with the maximal p-value for each stage is chosen to serve as an initial configuration for later simulations. p-value is calculated using a GLMM with Poisson distribution. The number of cells for the chosen time points ranges from 500 to 550 (the initial hexagonal arrays had exactly 400 cells). **(c)** Average histograms, comparing model's and experimental number of neighbors distributions. Each bar height is the average percentage of cells with a given number of neighbors. Scattered points are results from individual chosen initial configurations (green for E17.5 and orange for P0), or the results from individual live imaging experiments (cyan for E17.5 and pink for P0). Error bars show standard error of the mean.

The selected morphologies were subsequently used in two ways. In the lateral inhibition only model, cell shapes, contact lengths, and neighbor relationships were held constant throughout the simulation. In the hybrid model, the selected morphologies served as the initial tissue configurations and were allowed to evolve dynamically.

#### 5. Quantitative analysis pipeline for comparing simulated differentiation dynamics with live imaging measurements

We simulated the dynamics of Notch, Delta, and repressor levels on the 10 developmental stage specific morphologies for each developmental stage, generated as described in section 4. Simulations were initialized with near zero levels of Notch, Delta, and the repressor in all cells. The initial value of each variable in each cell was independently sampled from a uniform distribution between 0 and 0.01. The system of ordinary differential equations defined by Eqs. 7–9 was integrated using scipy.integrate.solve_ivp[4] with the RK45 method[5,6].

The differentiation dynamics obtained from the simulations are not directly comparable to those measured in the live imaging experiments. The main differences arise from three aspects of the model and experimental data: the classification of cells as HCs or SCs, the developmental state at the beginning of the observation period, and the absence of irreversible commitment to HC fate in the model. We therefore developed an analysis pipeline that accounts for each of these differences before comparing simulated and experimental differentiation statistics.

##### 5.1 Parameter-combination-specific Delta thresholds define simulated HC and SC populations

A key challenge was defining a Delta threshold, $\tilde{p}_D$, that separates HCs from SCs across different parameter combinations of $\tilde{p}_R$, $\tilde{p}_S$ and developmental stage (E17.5/P0). This threshold was used to classify every cell as HC or SC for the later analyses, i.e. a cell-$i$ is an HC if $\tilde{D}_i > \tilde{p}_D$ and a SC otherwise. The inclusion of contact length dependent signaling reduced the separation between the high-Delta/low-repressor and low-Delta/high-repressor states, resulting in a more continuous distribution of Delta and repressor levels. Consequently, a single fixed threshold cannot reliably distinguish differentiated from undifferentiated cells across all parameter combinations.

To determine an appropriate $\tilde{p}_D$ for each parameter combination $\left(\tilde{p}_R, \tilde{p}_S\right)$ and developmental stage, we performed 10 simulations for each parameter combination on each of the 10 corresponding developmental stage specific morphologies. Each simulation was initialized with different random initial conditions, resulting in 100 simulations per parameter combination and developmental stage.

For each simulation, the Delta levels at the final time point were sorted, and the largest gap in the distribution was identified. The minimum Delta value above this gap defined an upper bound, and the maximum Delta value below the gap defined a lower bound (Supplementary Figure 3a).

For each morphology, we then computed the maximum upper bound and the minimum lower bound across the 10 simulations, and defined the morphology-specific $\tilde{p}_D$ as the average of these two values (Supplementary Figure 3a). The final Delta threshold for each parameter combination and developmental stage was obtained by averaging the morphology-specific thresholds across the 10 morphologies (Supplementary Figure 3b), with the associated uncertainty given by the standard error of the mean (SEM).

The resulting threshold was used to classify cells as HCs or SCs in all simulations with the same parameter combination $\left(\tilde{p}_R, \tilde{p}_S\right)$ and developmental stage. The same thresholds were used for the hybrid model (Eq. 16-17).

#### 5.2 Edge statistics identify the simulation frame corresponding to the experimental first frame

Using the Delta threshold described in Section 5.1, we converted the simulated cell states at 100 sampled time points into binary HC/SC patterns. Each time point was compared with the initial experimental patterns from the corresponding developmental stage: simulations performed on E17.5-specific morphologies were compared with the E17.5 experimental data, whereas simulations performed on P0-specific morphologies were compared with the P0 data.

For each simulated time point, we calculated the fractions of HC:HC and HC:SC edges among all cell–cell interfaces: $p_{HC:HC} = \frac{N_{HC:HC}}{N_{edges}}, \quad p_{HC:SC} = \frac{N_{HC:SC}}{N_{edges}}$.

We then compared these fractions with the corresponding experimental values using the following objective function:

$$J_{initial}(\tilde{t}) = \frac{\left(p^{model}_{HC:HC} - \bar{p}^{exp}_{HC:HC}\right)^2}{\left(SEM^{exp}_{p_{HC:HC}}\right)^2} + \frac{\left(p^{model}_{HC:SC} - \bar{p}^{exp}_{HC:SC}\right)^2}{\left(SEM^{exp}_{p_{HC:SC}}\right)^2}. \qquad (24)$$

The experimental means and SEMs were calculated across the three biological replicates available for each developmental stage. The SC–SC edge fraction was not included because the three edge fractions satisfy $p_{HC:HC} + p_{HC:SC} + p_{SC:SC} = 1$ and therefore the SC–SC fraction contains no additional independent information.

Edge statistics were used for this alignment because HC-neighbor statistics alone tend to favor simulated frames containing very few HCs, even when the overall cell-type composition differs substantially from that of the experimental first frame. By contrast, edge fractions constrain both the relative abundance of HCs and SCs and their spatial arrangement.

For each simulation, the time point with the lowest $J_{initial}(\tilde{t})$ was selected as the 'chosen initial time-point', best matching the experimental initial pattern (Supplementary Figure 3c). Subsequent differentiation events were quantified relative to this matched frame, as described in Section 5.3.

#### 5.3 Differentiation events were defined relative to the first frame and steady state

In the live imaging experiments, differentiation events were identified directly from the emergence of Atoh1 expression. In the simulations, an SC-to-HC threshold crossing was counted as a differentiation event only if two conditions were satisfied: the cell was classified as an SC in the matched initial frame identified in Section 5.2, and it was classified as an HC in the final steady state of the simulation.

Because the model does not impose irreversible HC fate commitment, a cell could cross the Delta threshold multiple times during a simulation. In such cases, the last upward crossing of the threshold after the matched initial frame was defined as the differentiation event. The number of HC neighbors was evaluated at the corresponding simulation time point.

##### 5.4 Averaging across repeats accounts for initial-levels variability and averaging across morphologies accounts for tissue-level variability

To compare differentiation patterns across simulations with different numbers of cells and differentiation events, we normalized the number of newly differentiated HCs in each neighbor category by the total number of newly differentiated HCs. Cells were divided into three groups according to the number of HC neighbors present at their differentiation time: 0, 1 or ≥ 2. We defined:

$$p_k^{diff} = \frac{N_{diff,k}}{N_{diff}}, \qquad k \in \{0, 1, \geq 2\},$$

where $N_{diff,k}$ is the number of newly differentiated HCs with k HC neighbors at their differentiation time, and $N_{diff}$ is the total number of newly differentiated HCs. By definition, $\sum_k p_k^{diff} = 1$.

For each parameter combination $\left(\widetilde{p}_R, \widetilde{p}_S\right)$ and morphology, we calculated the mean and SEM of $p_k^{diff}$ across the 10 simulations initialized with different Notch, Delta, and repressor levels. In addition, we calculated the averaged $p_{HC:HC}^{model}$ and the average $p_{HC:SC}^{model}$ across the chosen initial time-point of each simulation. This first averaging step captures variability arising from the random initial signaling conditions. We then averaged the resulting morphology-level means across the 10 morphologies corresponding to each developmental stage. The SEM across these morphology-level means captures variability associated with tissue morphology.

##### 5.5 A combined objective function quantifies agreement between model and experiment

To quantitatively compare the model's and experimental results, we first re-calculated the $J_{initial}$, as a function of lateral inhibition sensitivity parameters, using the averaged $p_{HC:HC}^{model}$ and $p_{HC:SC}^{model}$ values:

$$J_{initial}(\tilde{p}_S, \tilde{p}_R) = \frac{\left(\bar{p}^{model}_{HC:HC} - \bar{p}^{exp}_{HC:HC}\right)^2}{\left(SEM^{exp}_{\bar{p}_{HC:HC}}\right)^2 + \left(SEM^{model}_{\bar{p}_{HC:HC}}\right)^2} + \frac{\left(\bar{p}^{model}_{HC:SC} - \bar{p}^{exp}_{HC:SC}\right)^2}{\left(SEM^{exp}_{\bar{p}_{HC:SC}}\right)^2 + \left(SEM^{model}_{\bar{p}_{HC:SC}}\right)^2}. \tag{25}$$

In addition, we compared the model and experimental differentiation statistics using the following objective function:

$$J_{diff}(\tilde{p}_S, \tilde{p}_R) = \sum_{k \in \{0,1\}} \frac{\left(\bar{p}^{diff,\, model}_{k} - \bar{p}^{diff,\, exp}_{k}\right)^2}{\left(SEM^{model}_{\bar{p}^{diff}_{k}}\right)^2 + \left(SEM^{exp}_{\bar{p}^{diff}_{k}}\right)^2}, \tag{26}$$

and define the combined objective function

$$J_{patterns}(\tilde{p}_S, \tilde{p}_R) = J_{initial}(\tilde{p}_S, \tilde{p}_R) + J_{diff}(\tilde{p}_S, \tilde{p}_R). \tag{27}$$

The model means and SEMs used in Eq. 25-26 were obtained after the two-step averaging procedure described in Section 5.4: first across simulation repeats and then across morphologies. The experimental means and SEMs were calculated across the three biological replicates.

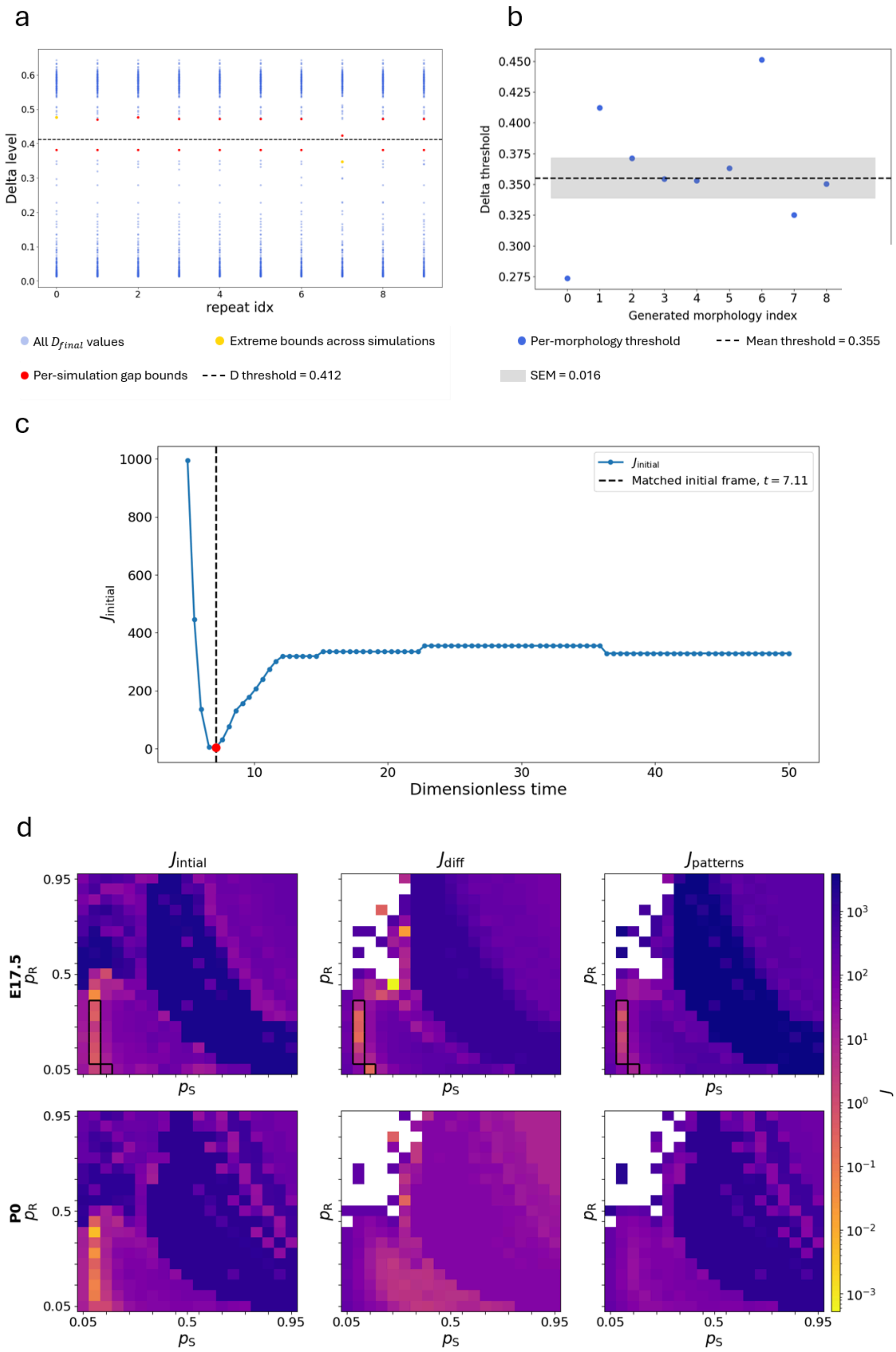


**Supplementary Figure 3. Quantitative analysis pipeline for comparing simulated differentiation dynamics with live imaging measurements**. **(a)** Construction of a Delta threshold for a single morphology at a representative parameter combination $\widetilde{p}_S = 0.1, \widetilde{p}_R = 0.3$. Each column shows the sorted final Delta levels of all cells in one simulation, with 10 simulations initialized using different signaling levels. Red points indicate the lower and upper bounds of the largest Delta gap identified in each simulation. Gold points indicate the minimum lower bound and maximum upper bound across simulations. The morphology-specific threshold was defined as the midpoint between these two values and is indicated by the dashed line. **(b)** Example of Delta-threshold averaging across the 10 generated E17.5 morphologies for a representative parameter combination $\tilde{p}_S = 0.1, \tilde{p}_R = 0.3$. Each point represents the morphology-specific threshold, the dashed line indicates the mean across morphologies, and the shaded region denotes the SEM. The same averaging procedure was applied independently to every parameter combination and to both developmental stages. This averaged threshold is used to classify HCs and SCs for all simulations with the same parameter combination and developmental stage.

**(c)** Objective function $J_{initial}(\tilde{t})$ as a function of dimensionless simulation time (Eq. 24) for an E17.5-specific morphology with $\tilde{p}_S = 0.1$ and $\tilde{p}_R = 0.3$. The score quantifies the agreement between the simulated and experimental HC:HC and HC:SC edge fractions in the initial imaging frame. The simulation time point with the minimum $J_{initial}(\tilde{t})$ was selected as the frame corresponding to the experimental initial pattern. The dashed line and red marker indicate the selected frame at $t = 7.11$, for which $J_{initial} = 3.41$.

**(d)** Objective functions scores across the $\left(\tilde{p}_S, \tilde{p}_R\right)$ parameter space. The left column shows the initial frame edge score, $J_{initial}$; the middle column shows the differentiation-time HC-neighbor score normalized by the total number of newly differentiated HCs, $J_{diff}$; and the right column shows $J_{patterns}$. E17.5 results are shown in the upper row and P0 results in the lower row. Lower values indicate better agreement between model and experiment. For $J_{diff}$, and subsequently $J_{patterns}$, white regions indicate undefined scores. For $J_{diff}$, undefined values occur when the matched initial frame already contains all HCs present in the final state, leaving no subsequent differentiation events and therefore $N_{diff} = 0$. Several parameter combinations satisfied $J_{patterns} < 16$ at E17.5 (marked by black rectangles), whereas none did so at P0.

#### 6. Contact-dependent lateral inhibition model with fixed morphologies reproduces E17.5 but not P0 differentiation statistics

Using the objective functions defined in Eqs. (25-27), we searched for parameter combinations for which the lateral inhibition model reproduced the experimental HC pattern and differentiation statistics at E17.5 and P0. Our search is done for $p_R$ and $p_S$ values in the range 0.05-0.95, with 0.05 resolution. Qualitatively, the model generated HCs that differentiated adjacent to existing HCs, as observed experimentally, rather than producing only an ideal salt-and-pepper pattern (Main Text Figure 3b).

We used the combined objective function $J_{patterns}(\tilde{p}_S, \tilde{p}_R)$ to fit the lateral inhibition sensitivity parameters. As this function includes four independent $\chi^2$ terms, we considered parameter combinations with $J_{patterns}(\tilde{p}_S, \tilde{p}_R) < 16$ (an average distance of less than two combined SEMs from the mean) to provide a good match (Supplementary Figure 3d).

Results indicated that the model reproduced the E17.5 data substantially better than the P0 data. The minimal $J_{patterns}(\tilde{p}_S, \tilde{p}_R)$ was 0.671 for E17.5, compared with 25.290 for P0 (Main

Text Figure 3c-d). Thus, several parameter combinations satisfied the selected fitting criterion for E17.5, whereas no P0 parameter combination did. These results indicate that contact length dependent signaling introduces sufficient variability in cell-cell communication to reproduce the differentiation of HCs next to existing HCs to quantitatively fit the E17.5 result (Main Text Figure 3c-d). However, the low extent of these HC differentiation next to existing HCs, and the lack of SCs with no HC neighbors (Extended Data Figure 6d) prevented the model from fitting the P0 differentiation patterns (Main Text Figure 3c-d).

#### 7. Fitting 2D vertex model's mechanical parameters

With the fitted lateral inhibition sensitivity parameters in hand, we turned to fitting the mechanical parameters. For this, we chose a few observables to compare between model and experimental results: (i) As a circle is the shape with the smallest perimeter for a given area, we anticipate roundness to be controlled by the contractility, we therefore used the HCs to SCs cell roundness ratio as the first observable; (ii) Area elasticity is anticipated to control how much a cell would change its area when one of its neighbors delaminate, we therefore used the HCs to SCs area change next to ablation ratio as the second observable; (iii) as our mechanical properties measurement relays of shrinking after circular ablation (Main Text Figure 4), we used the linear shrinking as the third observable; (iv) The final observable is the mechanical stress, which is not dimensionless, and so cannot be compared directly to the model results. Instead, we compared the mechanical stress ratio between E17.5 and P0. For the differentiation threshold $\tilde{p}_D$ in Eq. 16-17, we used the threshold determined (see section 5.1 above) for the E17.5 lateral inhibition parameters. The sections below explain in detail how each observable was calculated for the model results and how it was compared with the experimental results.

##### 7.1 HCs to SCs roundness ratio

The roundness of individual cells ($4\pi A_n / L_n^2$) was measured for the final frame of each live imaging control experiment (with no treatment). The HC to SC average roundness ratio was then calculated for each experiment (Extended Data Figure 6b). To fit simulation results to this data, the exact same calculation was applied for the final frame of each simulation and the roundness score is calculated according to

$$\chi^2_{roundness} = \frac{\left(<model\ roundness\ ratio> - <experimental\ roundness\ ratio>\right)^2}{\left(experimental\ roundness\ ratio\ SEM\right)^2}. \qquad (28)$$

##### 7.2 HCs to SCs area change next to ablation ratio

The area of all cells that were the direct neighbors of an ablated cell, for each single HC ablation experiment, was measured just before ablation and one hour after ablation (when the created space in the apical plane is filled by neighboring cells). Using tracking results, the after-ablation to before-ablation area ratio was calculated for each cell. Then, the average area ratio was calculated for all HCs and SCs at each experiment and the ratio of HCs area change to SCs area change is calculated. To fit simulation results to this data, ablation of four chosen cells was simulated over each array, after simulation has reached a steady-state and subsequently simulated until reaching a steady-state again. Then HCs to SCs area change ratio was calculated in the same way as done for experiments, using the before ablation steady-state to calculate the before areas and the after ablation steady-state to calculate the after areas. The ablation score was calculated according to

$$\chi^2_{abl-area} = \frac{\left(<model\ area\ change\ after\ ablation\ ratio> - <experimental\ area\ change\ after\ ablation\ ratio>\right)}{\left(experimental\ area\ change\ after\ ablation\ ratio\ \ SEM\right)^2}. \quad (29)$$

**7.3 Circular ablation shrinkage percentage**

To fit the circular ablation experiments (Main text Figure 4), we started by calculating the average radius shrinking, $\lambda_{exp} = \frac{R_{final}}{R_{initial}}$, for each stage. Assuming uniform isotropic shrinking, we estimated the linear shrinking for the model by scaling all distances by multiplying with a scaling parameter λ. This scales perimeters by λ and areas by $\lambda^2$. Plugging it into the model's energy we got

$$\tilde{E}(\lambda) = \sum_{n\in cells}\left[\frac{\tilde{\alpha}_n}{2}(\lambda^2\tilde{\tilde{A}}_n - \tilde{\tilde{A}}_0)^2 + \frac{\tilde{\Gamma}_n}{2}\lambda^2\tilde{\tilde{L}}_n^2\right]. \quad (30)$$

We minimized this energy with respect to λ by solving $\frac{\partial\tilde{E}}{\partial\lambda} = 0$ to get the predicted shrinking:

$$\frac{\partial\tilde{E}}{\partial\lambda} = 0 \Rightarrow \lambda^2 = 0\ or\ \lambda_{model}^2,\ where\ \lambda_{model}^2 \equiv \frac{\sum_{n\in cells}\left[2\tilde{\tilde{A}}_0\tilde{\tilde{A}}_n\tilde{\alpha}_n - \tilde{\Gamma}_n\tilde{\tilde{L}}_n^2\right]}{2\sum_{n\in cells}\tilde{\tilde{A}}_n^2\tilde{\alpha}_n}. \quad (31)$$

For a non-negative shrinking, $\lambda = 0$ maximizes the mechanical energy while $\lambda_{model}$ minimizes it.

Given a set of steady-state morphologies and Delta levels from model simulations, we could plug in the specific values to Eq. 31 and find $\lambda_{model}$ for each, then compare it to the measured experimental value by

$$\chi^2_{shrinkage} = \frac{\left(<\lambda_{model}> - <\lambda_{exp}>\right)^2}{\left(\lambda_{exp}SEM\right)^2}. \quad (32)$$

To estimate the needed $\tilde{A}_0$ for a good fit one can substitute $\lambda_{model}$ by $\lambda_{exp}$ in Eq. 31, which gives the following preferred area:

$$\tilde{\tilde{A}}_0 = \frac{\sum_{n\in cells} \tilde{\Gamma}_n \tilde{\tilde{L}}_n^2 + 2\lambda_{exp}^2 \sum_{n\in cells} \tilde{\tilde{A}}_n^2 \tilde{\alpha}_n}{2 \sum_{n\in cells} \tilde{\tilde{A}}_n \tilde{\alpha}_n} . \quad (33)$$

As an initial guess one can calculate $\tilde{\tilde{A}}_0$ for an isotropic tissue where all cells have the same circular shape diameter $a$, i.e. $\tilde{\tilde{A}}_n = \frac{\pi}{4}$, $\tilde{\tilde{L}}_n = \pi$. With this approximation we got

$$\tilde{\tilde{A}}_0 \approx 2\pi \frac{\sum_{n\in cells} \tilde{\Gamma}_n}{\sum_{n\in cells} \tilde{\alpha}_n} + \frac{\pi}{4}\lambda_{exp}^2 . \quad (34)$$

##### 7.4 Mechanical stress ratio

The model mechanical stress was calculated in a similar way to linear shrinkage (for the case with uniform areas and perimeters):

$$\sigma = \frac{\partial \tilde{E}}{\partial \tilde{\tilde{A}}_{tot}} = \sum_{n\in cells} \frac{\tilde{\tilde{A}}_n}{\tilde{\tilde{A}}_{tot}} \frac{\partial \tilde{E}}{\partial \tilde{\tilde{A}}_n} = \sum_{n\in cells} \frac{\tilde{\tilde{A}}_n}{\tilde{\tilde{A}}_{tot}} \left[ \tilde{\alpha}_n (\tilde{\tilde{A}}_n - \tilde{A}_0) + \tilde{\Gamma}_n \tilde{\tilde{L}}_n \frac{\partial \tilde{\tilde{L}}_n}{\partial \tilde{\tilde{A}}_n} \right] . \quad (35)$$

Approximating the cells as circles with the same areas and substituting $\tilde{\tilde{A}}_0$ for the approximated value from (Eq. 34) we got

$$\sigma \approx \frac{\pi}{4}(1 + \lambda_{exp}^2) \frac{1}{N} \sum_{n\in cells} \tilde{\alpha}_n \approx \frac{\pi}{4}(1 + \lambda_{exp}^2)(f_{HC}\alpha_{HC} + f_{SC}\alpha_{SC}), \quad (36)$$

where $f_{HC}$ is the fraction of HCs and $f_{SC}$ is the fraction of SCs in the array. We used the perimeter-area relations for a circle $L = 2\sqrt{A\pi} \Rightarrow \frac{\partial L}{\partial A} = \sqrt{\frac{\pi}{A}} = 2$. This can be used to derive the ratio between $\alpha_{HC}$ for the different stages according to the ratio of measured mechanical stress for each stage:

$$\frac{(1+\lambda_{exp}^{P0^2})\ (f_{HC}^{P0}\alpha_{HC}^{P0}+f_{SC}^{P0})}{(1+\lambda_{exp}^{E17.5^2})\ (f_{HC}^{E17.5}\alpha_{HC}^{E17.5}+f_{SC}^{E17.5})} = \frac{\tilde{\sigma}_{exp}^{P0}}{\tilde{\sigma}_{exp}^{E17.5}}. \quad (37)$$

##### 7.5 Finding the model's mechanical parameters

There are 4 free fitting parameters in the 2D vertex model - $\tilde{\alpha}_{HC}$, $\tilde{\Gamma}_{SC}$, $\tilde{\Gamma}_{HC}$ and $\tilde{\tilde{A}}_0$. These parameters are fitted for E17.5 and P0 separately, minimizing the combined objective function

$$J_{mech}(\tilde{\tilde{A}}_0, \tilde{\alpha}_{HC}, \tilde{\Gamma}_{SC}, \tilde{\Gamma}_{HC}) = \chi^2_{roundness} + \chi^2_{abl-area} + \chi^2_{shrinkage}. \quad (38)$$

For each combination ($\tilde{\alpha}_{HC}$, $\tilde{\Gamma}_{SC}$, $\tilde{\Gamma}_{HC}$) values, we calculated this objective function via the following steps:

1. We started with the best matching steady-state patterns, produced by the LI only model for each initial disordered configuration (i.e. the ones with the lowest $J_{patterns}$ ). The selected threshold for each pattern is then used to determine the mechanical properties of each cell, by setting the $\tilde{p}_D$ value in Eq. 16-17..
2. We calculated the initial guess for $\tilde{\tilde{A}}_0$ according to Eq. 34.
3. We ran a mechanical simulation only (no differentiation) until reaching a steady state.
4. We calculated $\chi^2_{roundness}$ and $\chi^2_{shrinkage}$ according to Eq. 28 and 32 respectively.
5. We updated $\tilde{\tilde{A}}_0$ value according to Eq. 33 using the steady-state perimeters and areas and reran the simulation until $\tilde{\tilde{A}}_0$ value converges. This convergence had been achieved in 2-4 iterations for every parameters combinations, resulting in a decrease in $\chi^2_{shrinkage}$ without increasing $\chi^2_{roundness}$.
6. We simulated the ablation of 4 chosen cells (same cells for each simulation) and calculated $\chi^2_{abl-area}$ according to Eq. 29.

The search through the 3D parameter space spanned by ($\tilde{\alpha}_{HC}$, $\tilde{\Gamma}_{SC}$, $\tilde{\Gamma}_{HC}$) was done via the following steps:

1. First, we fitted E17.5 experimental results. We started with the simplifying assumption that $R_{HC:SC} \equiv \frac{\tilde{\alpha}_{HC}}{\tilde{\alpha}_{SC}} = \frac{\tilde{\Gamma}_{HC}}{\tilde{\Gamma}_{SC}}$, which reduced the parameter space to 2D. We also limited the search for regions where $R_{HC:SC}$>1, due to the experimental

observation that HCs are rounder than SCs ($\frac{\tilde{\Gamma}_{HC}}{\tilde{\Gamma}_{SC}}$>1) and change their areas less near ablation ($\frac{\tilde{\alpha}_{HC}}{\tilde{\alpha}_{SC}}$>1). In addition, we limited $\tilde{A}_0$ to be smaller than the average cell area, which is roughly $\pi/4$. According to Eq. 34, together with the simplifying assumption, this condition on $\tilde{A}_0$ translate into the condition $\tilde{\Gamma}_{SC} < \frac{1+\lambda_{exp}^{2}}{8} \approx 0.018$. For this 2D search we used $R_{HC:SC} \in \{1.25,\ 1.75, 2.5,\ 3.5,\ 5.0\}$ (i.e geometric series with ratio $\sim\sqrt{2}$) and $\tilde{\Gamma}_{SC} \in \{0.002,\ 0.006,\ 0.010,\ 0.014,\ 0.0175\}$ (i.e. linear series with $0.004$ spacing). The 2D search results yielded a good fit for the E17.5 results with $\chi^2_{mech} = 4.1$ for $R_{HC:SC} = 3.5, \tilde{\Gamma}_{SC} = 0.0175$. In addition, results showed that the roundness score is sensitive to $R_{HC:SC}$, while essentially not at all to $\tilde{\Gamma}_{SC}$ (Supplementary figure 4a).

2. We improved the fit by fixing $\tilde{\alpha}_{HC} = 3.5$ and $\tilde{\Gamma}_{SC} = 0.0175$ and relaxing the $\frac{\tilde{\alpha}_{HC}}{\tilde{\alpha}_{SC}} = \frac{\tilde{\Gamma}_{HC}}{\tilde{\Gamma}_{SC}}$ assumption. We then performed a 1D search for $\frac{\tilde{\Gamma}_{HC}}{\tilde{\Gamma}_{SC}} \in \{1.1,\ 1.25,\ 1.4,\ 1.55,\ 1.757,\ 1.85,\ 1.94\}$ (i.e. linear series with spacing of 0.15, but including the previously existing point at 1.757). The upper bound was given by the highest $\tilde{\Gamma}_{HC}$ value giving $\tilde{\tilde{A}}_0 < \frac{\pi}{4}$ according to Eq. 34 with the given $\tilde{\Gamma}_{SC}$ and $\tilde{\alpha}_{HC}$ values.
3. As the best result yet has been achieved for the upper bound $\frac{\tilde{\Gamma}_{HC}}{\tilde{\Gamma}_{SC}}$ value, we performed another search for $\tilde{\Gamma}_{SC} \in \{0.009,\ 0.0105,\ 0.0125,\ 0.0145,\ 0.0175\}$ (i.e. geometric series with a ratio of ~$\sqrt{2}$), setting $\tilde{\Gamma}_{HC}$ to its upper bound for each $\tilde{\Gamma}_{SC}$ value, according to Eq. 34 with $\tilde{\tilde{A}}_0 = \frac{\pi}{4}$ (Supplementary figure 4b left). This gave the best fit for $\tilde{\alpha}_{HC} = 3.5, \tilde{\Gamma}_{SC} = 0.0105, \frac{\tilde{\Gamma}_{HC}}{\tilde{\Gamma}_{SC}} = 6.67$ and $\tilde{\tilde{A}}_0 = 0.74181$ with $J_{mech} = 1.242$.
4. For fitting P0 results we first set the $\tilde{\alpha}_{HC}$ parameter using the E17.5 results, according to Eq. 37:

$$\alpha_{HC}^{P0} = \frac{\tilde{\sigma}_{exp}^{P0}\,(1+\lambda_{exp}^{E17.5^2})\,(f_{HC}^{E17.5}\alpha_{HC}^{E17.5}+f_{SC}^{E17.5})}{\tilde{\sigma}_{exp}^{E17.5}\,(1+\lambda_{exp}^{P0^2})\,f_{HC}^{P0}} - \frac{f_{SC}^{P0}}{f_{HC}^{P0}} \approx 1.757, \qquad (39)$$

using the average experimental values as $\lambda_{exp}$ for each stage, the average number of HCs and SCs in the initial arrays for the $N_{SC}$ and $N_{HC}$, and the ratio between the average measured mechanical stresses for E17.5 and P0 (see main text Figure 4d).

5. We repeated step 3 with the calculated P0 $\tilde{\alpha}_{HC}$ value, i.e. performed a search for $\tilde{\Gamma}_{SC} \in \{0.0075,\ 0.009,\ 0.0105,\ 0.0125,\ 0.0145\}$ (i.e. geometric series with a ratio of ~$\sqrt{2}$), setting $\tilde{\Gamma}_{HC}$ to its upper bound, according to Eq. 34 with $\tilde{A}_0 = \frac{\pi}{4}$ (Supplementary figure 4b right). This yielded the best fitting score for $\tilde{\alpha}_{HC} = 1.757,\ \tilde{\Gamma}_{SC} = 0.0105, \frac{\tilde{\Gamma}_{HC}}{\tilde{\Gamma}_{SC}} = 4.09, \tilde{\tilde{A}}_0 = 0.7585$ with $J_{mech} = 9.837$.

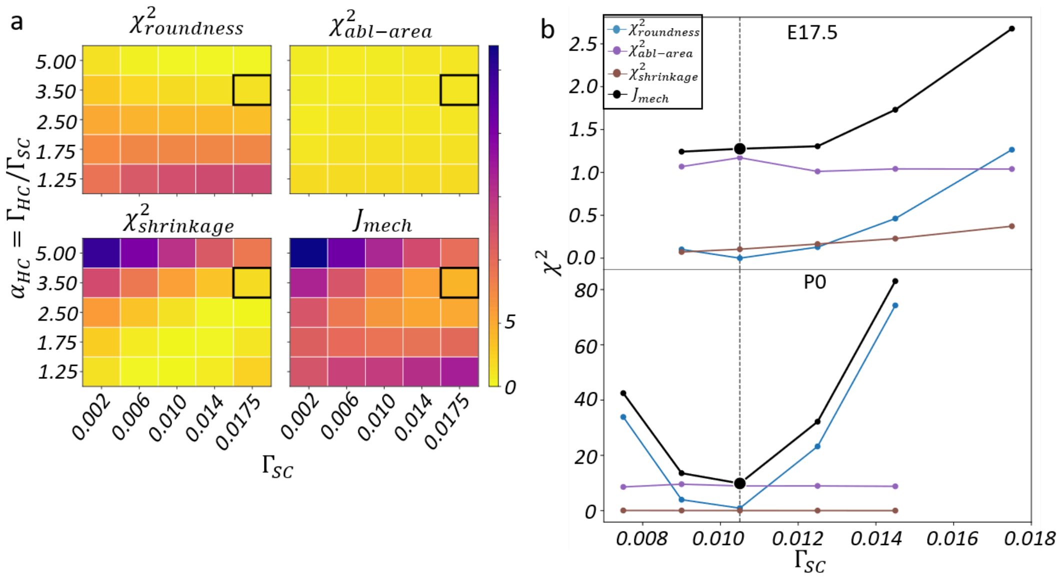


**Supplementary Figure 4. Fitting mechanical parameters. (a)** E17.5 mechanical parameters are first fitted under the simplifying assumption $\tilde{\alpha}_{HC} = \tilde{\Gamma}_{HC}/\tilde{\Gamma}_{SC}$, performing a 2D search over different $(\tilde{\Gamma}, \tilde{\alpha}_{HC})$ pairs and their corresponding $\tilde{\tilde{A}}_0$ value. For each pair, simulation results are compared with experimental roundness ratio (top left), experimental area change near ablation (top right) and experimental shrinkage after circular ablation (bottom left), to get the summed objective function $J_{mech}$ (bottom right). Results show that $\chi^2_{roundness}$ is not sensitive to $\tilde{\Gamma}_{SC}$ (top left). The best fit (marked by black rectangles) was achieved for the upper boundary of $\tilde{\Gamma}_{SC}$. This search sets the value of $\tilde{\alpha}_{HC}$ for E17.5 while the value for P0 is calculated based on the ratio between the experimentally measured mechanical stress for both stages. **(b)** To fit $\tilde{\Gamma}_{SC}$ and $\tilde{\Gamma}_{HC}$ for each stage, a search is made over $\tilde{\Gamma}_{SC}$ values where $\tilde{\Gamma}_{HC}$ is set to the highest possible value that still gives $\tilde{\tilde{A}}_0 < \pi/4$ according to Eq. 34. The lowest $J_{mech}$ value (black curve, best point marked with a vertical line) is achieved for the same $\tilde{\Gamma}_{SC}$ value for both stages (resulting in a different $\tilde{\Gamma}_{HC}$ value as the upper boundary depends on $\tilde{\alpha}_{HC}$).

One point worth noting - for the experimental results, on average, HCs tend to decrease their apical area after one of their neighboring cells was ablated. This effect could not be reproduced by the model, where cells always increase their area when a neighbor is ablated. Therefore, the large $\chi^2_{abl-area}$ for P0 is a real model limitation. The same limitation applies also to E17.5, but there the experimental standard error is much larger, resulting in a good $\chi^2_{abl-area}$ score.

Final fit results are summarized in Extended Data Figure 6b.

#### 8. Hybrid model

To combine the lateral inhibition differentiation model with the 2D vertex mechanical model we implemented a hybrid model. In this model, Notch, Delta and Repressor levels for each cell are updated according to Eq. 7,9,19 in parallel to the 2D vertex model simulation, which includes minimizing the mechanical energy (Eq. 14-15). This is done in a quasi-static fashion, due to the observed fast mechanical response (~a few minutes, see circular ablation results at Main text Figure 4) compared with slow differentiation (~30 hours, see differentiation after single HC ablation, main text Figure 2c-d). The 2D vertex model is simulated until reaching a steady state, then lateral inhibition levels (Notch, Delta and Repressor) are updated for the entire time interval. In this model, Lateral inhibition and tissue mechanics influence each other in three ways: (i) contact lengths are updated according to tissue mechanics, affecting lateral inhibition dynamics (Eq. 7,9,19 through the $\tilde{S}_n$ and $\tilde{G}_n$ terms); (ii) the mechanical parameters of each cell ($\tilde{\alpha}_n$, $\tilde{\Gamma}_n$) are updated according to its Delta level (Eq. 16-17); (iii) In the version with mechanosensitivity, changing mechanical tension affects Delta production directly. For this model we used the E17.5 fitted lateral inhibition parameters $\tilde{p}_S$, $\tilde{p}_R$ and their corresponding threshold $\tilde{p}_D$, both for E17.5 and P0 simulations. In addition, we used the stage specific initial disordered configurations and fitted mechanical parameters.

##### 8.1 Comparing hybrid-model and experimental differentiation patterns

To compare the experimental differentiation pattern with the hybrid model results we used the exact same scheme as for the lateral inhibition only model (see section 5 above), using the $J_{patterns}$ score (Eq. 27), but this time as a function of $\tilde{p}_T$.

$$J_{patterns}(\tilde{p}_T) = J_{initial}(\tilde{p}_T) + J_{diff}(\tilde{p}_T). \tag{40}$$

Starting with no mechanosensitivity, $\tilde{p}_T = 0$, we performed three hybrid model simulations on each of the 10 corresponding developmental stage specific morphologies (same as for the lateral inhibition only model), for each stage, until reaching a steady state. $J_{patterns}(\tilde{p}_T)$ was calculated for these simulations (Main text Figure 6b-c).

**8.2 Estimating required mechanosensitivity**

Next, we used these results and calculated the average $\tilde{Y}_n$ levels (Eq. 23) as a function of time along the simulation, according to cell type and the number of HC neighbors, averaging over repeats first, then using these averages to calculate the average and SEM over morphologies. To be clear, these $\tilde{Y}_n$ levels had no effect on the no-mechanosensitivity simulation. Results showed a clear distinction between the groups. HCs had the largest $\tilde{Y}_n$ levels for most of the simulation time (after the initial patterning phase, where all cells are SCs), followed by SCs with two or more HC neighbors (Supplementary Figure 5a). The group of SCs with no HC neighbors started with the lowest $\tilde{Y}_n$ levels and later swapped places with the SCs with one HC neighbor group, which started as the second lowest (Supplementary Figure 5a). Comparing stages, the $\tilde{Y}_n$ trends were similar, but the E17.5 cells had overall larger $\tilde{Y}_n$ levels than the P0 cells (Supplementary Figure 5a). According to these results, we estimated that $\tilde{p}_T$ values between 0.06 and 0.12 would have the strongest discriminating effect.

**8.3 Fitting mechanosensitivity**

The mechanosensitivity threshold $\tilde{p}_T$ was assumed to be the same for both stages and fitted jointly. Since $\tilde{p}_T = \frac{p_T}{\alpha_{SC} a^3 L_0}$ joint $\tilde{p}_T$assumption implies that, besides the same $\tilde{p}_T$, the SC area elasticity $\alpha_{SC}$ and the length scales $a$ and $L_0$ are the same at E17.5 and P0. We ran the hybrid model simulation with $\tilde{p}_T$ between 0.06 and 0.12 with 0.01 interval, scoring these runs, we found that between $\tilde{p}_T = 0.08$ gave the best results for the sum of $J_{patterns}(\tilde{p}_T)$ over E17.5 and P0 (Supplementary Figure 5b), for $\tilde{p}_T \geq 0.1$ more than 50% of the simulations did not produce any pattern (characterized by the percentage of HCs below 10% at steady-state). We then ran a finer grid search for $\tilde{p}_T$ between 0.07 and 0.09 with 0.033 interval, finding the best joint score at $\tilde{p}_T = 0.0866$, followed by an ultra-fine grid search for $\tilde{p}_T$ between 0.0833 and 0.088 with 0.0011 interval yielding the best fitting point for $\tilde{p}_T = 0.0855$ (Supplementary Figure 5b inset). Note that although $\tilde{p}_T = 0.0866$ yielded a slightly better $J_{patterns}(\tilde{p}_T)$ sum over stages (~18 compared to ~20 for $\tilde{p}_T = 0.0866$), it had far less differentiation events at P0 (13 compared to 66 for $\tilde{p}_T = 0.0866$). As a consequence, the lower $J_{patterns}(\tilde{p}_T)$ for P0 with $\tilde{p}_T = 0.0866$, stemmed mostly from high SEM of the model results. We therefore preferred the $\tilde{p}_T = 0.0855$ result with slightly higher $J_{patterns}(\tilde{p}_T)$ but lower SEMs for P0 (~10 compared to ~25 for $\tilde{p}_T = 0.0866$).

To validate the mechanosensitive model with the fitted $\tilde{p}_T$ value, we calculated the percentage of isolated SCs for the hybrid model results, without mechanosensitivity and for $\tilde{p}_T = 0.0855$. With mechanosensitivity we got many more isolated SCs for the P0 fitted mechanical parameters, in qualitative agreement with the experimental results (Extended Data Figure 6d).

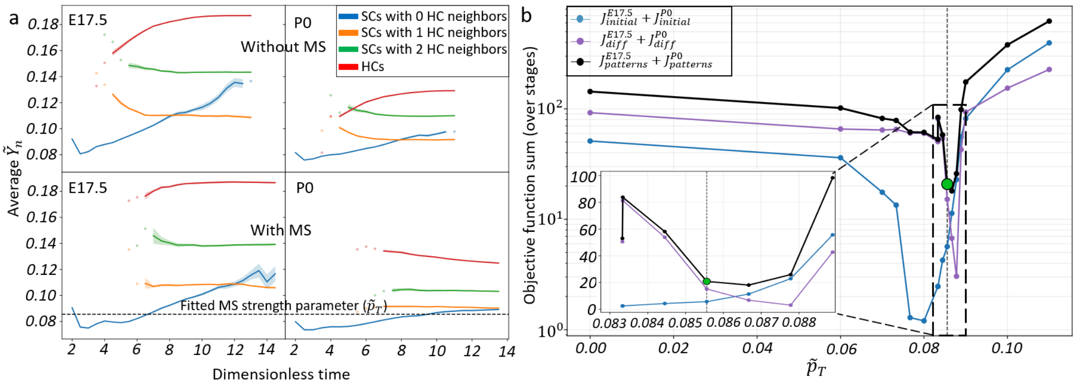


**Supplementary Figure 5. Fitting mechanosensitivity strength. (a)** Simulated average tension (summed over each cell's boundaries, averaged over all cells in each group) for the hybrid model (top/bottom – without/with mechanosensitivity, left/right E17.5/P0), grouped by cell type and its number of HC neighbors (for SCs). For most simulated time points, SCs with more HC neighbors have higher tension. Solid lines track group averages over all simulations (3 repeats for each of the 10 initial arrays for each stage and model), standard errors are shown as shaded areas around each line. Horizontal dashed line marks the chosen $\tilde{p}_T$ value. Scattered points are shown for time points where the group is not present in at least one repeat for each array (HCs take time to differentiate while SCs with no HC neighbors do not always exist at steady state). **(b)** Objective functions (summed over developmental stages) as a function of the mechanosensitivity strength parameter $\tilde{p}_T$ ($\tilde{p}_T$=0 means no mechanosensitivity) on a semi-log scale. The chosen value is marked (green circle and vertical line). Inset shows an enlarged, linear scale plot of the fine-resolution search region (marked by dashed rectangle). MS – Mechanosensitivity.

#### 8.4 Single HC ablation in the hybrid model

With the fitted $\tilde{p}_T$ value in hand, we sought to get a prediction for the distance of differentiation events from ablation at P0. For that we used the resulting steady-states, from the first repeat for each P0 initial morphology, chose a random HC and ablated it, and simulated for additional 5 time units after ablation. We simulated 5 repeats for each array, choosing a different HC to ablate at each repeat. Next, we detected differentiation events, defined as before, cells that were classified as SCs before ablation and turned to HCs by the end of the simulation. We then measured the distance (in cell perimeter units) between the ablated HC and the future differentiating cells, just before ablation (same as done for single HC ablation experiments, see main text Fig. 2e). To compare this distance with the experimentally measured distance, we converted the experimental results, measured in $\mu m$ to average cell diameter units. The average cell diameter for each experiments was calculated for the frame just before ablation (same frame for which the distance from ablation measurements were done), estimating it via the circular approximation, $diameter \approx 2\sqrt{area/\pi}$. Model results show a better quantitative agreement with experiments, compared to the model without mechanosensitivity (Main text Figure 6f).

### Supplementary Tables

| Parameter Name | Parameter Meaning | Reasoning | Value |
|---|---|---|---|
| $l$ | Hill functions exponent | Assuming cooperative binding | 3 |
| $L_0$ | The typical length scale | Assuming it is equal to the average cell perimeter per morphology | $\langle L_i \rangle$ |
| $\tau_N$, $\tau_D$ | Relative degradation rates for Notch and Delta to Repressor | Assuming equal degradation rate | 1 |
| $\tilde{\beta}$ | Ratio between the production rate of Notch and Delta | Assuming equal production rates of Notch and Delta | 1 |
| min_bond_length | Virtual vertices are removed for edges with smaller lengths | Chosen to be a bit larger than the intercalation threshold | 0.05 |
| max_bond_length | Virtual vertices are added for edges with larger lengths | Balancing running time and number of virtual vertices | 0.2 |
| intercalation_length | Edges smaller than this length intercalate | ~ 10% of the typical edge length | 0.04 |
| delamination_area | Cells with smaller area delaminate | ~ 10% of the typical cell area | 0.1 |
| division_area | Cells with larger area divide | ~150% of the typical cell area | 1.2 |
| sharp_angle_threshold | Vertices with smaller angles are removed | Preventing edge crossing | 0.1 |

| | | | |
|---|---|---|---|
| li_steady_state_threshold | Lateral inhibition steady-state is announced when all lateral inhibition derivatives are smaller than this threshold for at least 3 time steps | Based on measuring the steady-state fluctuation in the simulations | 0.01 |
| Initial_dt | Initial and maximal time interval | Largest with not many edge crossings | 0.01 |
| dt_decrease_factor | dt is multiplied by this factor when the vertex moved too far | Arbitrary value less than 1 | 0.5 |
| dt_increase_factor | dt is multiplied by this factor after a successful time step, if it is smaller than the initial_dt | Arbitrary value more than 1 | 1.1 |
| low_dt_threshold | dt cannot decrease below this value | Arbitrary small value | 0.0001 |
| max_displacement | A vertex cannot move more than this distance in a single time step, if it does, step is reverted and dt decreased | Quarter of the min_bond_length | 0.0125 |
| Mechanical steady-state threshold | A maximal vertex displacement less than this threshold for 3 consecutive steps would determine a mechanical steady-state | Noise analysis for manually verified steady-state | 0.003 |
| Lateral inhibition steady-state threshold | A maximal change in lateral inhibition levels less than this threshold for 3 consecutive steps would determine a lateral inhibition steady-state | Noise analysis for manually verified steady-state | 0.001 |

**Supplementary Table 1. Additional Model parameters.** Parameters used for model simulation, in addition to the fitted free parameters reported in Extended Data Table 1,2.

| Disordered array number | Number of cells at the best matching frame for E17.5 | Number of cells at the best matching frame for P0 | p-value of the best matching frame for E17.5 | p-value of the best matching frame for P0 |
|---|---|---|---|---|
| 0 | 508 | 509 | 0.9933 | 0.9972 |
| 1 | 537 | 538 | 0.9772 | 0.9900 |

| | | | | |
|---|---|---|---|---|
| 2 | 516 | 519 | 0.9948 | 0.9830 |
| 3 | 523 | 525 | 0.9996 | 0.9951 |
| 4 | 518 | 523 | 0.9837 | 0.9976 |
| 5 | 524 | 531 | 0.9963 | 0.9987 |
| 6 | 530 | 538 | 0.9957 | 0.9962 |
| 7 | 520 | 524 | 0.9904 | 0.9987 |
| 8 | 517 | 518 | 0.9945 | 0.9925 |
| 9 | 524 | 527 | 0.9822 | 0.9879 |

**Supplementary Table 2. Disordered arrays p-values.** p-values for the selected best matching randomly created patterns, comparing their number of neighbors distribution with the experimental one (from the first frame of each experiment). p-values were calculated using a generalized linear mixed model with Poisson distribution.
